# Conservation and evolution of the sporulation gene set in diverse members of the Firmicutes

**DOI:** 10.1101/2022.02.25.481979

**Authors:** Michael Y. Galperin, Natalya Yutin, Yuri I. Wolf, Roberto Vera Alvarez, Eugene V. Koonin

## Abstract

The current classification of the phylum Firmicutes (new name, *Bacillota*) features eight distinct classes, six of which include known spore-forming bacteria. In *Bacillus subtilis*, sporulation involves up to 500 genes, many of which do not have orthologs in other bacilli and/or clostridia. Previous studies identified about 60 sporulation genes of *B. subtilis* that were shared by all spore-forming members of the Firmicutes. These genes are referred to as the sporulation core or signature although many of these are found also in genomes of non-spore-formers. Using an expanded set of 180 firmicute genomes from 160 genera, including 76 spore-forming species, we investigated the conservation of the sporulation genes, in particular, seeking to identify lineages that lack some of the genes from the conserved sporulation core. The results of this analysis confirmed that many small acid-soluble spore proteins (SASPs), spore coat proteins, and germination proteins, which were previously characterized in bacilli, are missing in spore-forming members of *Clostridia* and other classes of Firmicutes. A particularly dramatic loss of sporulation genes was observed in the spore-forming members of the families *Planococcaceae* and *Erysipelotrichaceae*. Fifteen species from diverse lineages were found to carry *skin* (*sigK*-interrupting) elements of different sizes that all encoded SpoIVCA-like recombinases but did not share any other genes. Phylogenetic trees built from concatenated alignments of sporulation proteins and ribosomal proteins showed similar topology, indicating an early origin and subsequent vertical inheritance of the sporulation genes.

**IMPORTANCE:** Many members of the phylum Firmicutes (*Bacillota*) are capable of producing endospores, which enhance the survival of important Gram-positive pathogens that cause such diseases as anthrax, botulism, colitis, gas gangrene, and tetanus. We show that the core set of sporulation genes, defined previously through genome comparisons of several bacilli and clostridia, is conserved in a wide variety of spore-formers from several distinct lineages of Firmicutes. We also detect widespread loss of sporulation genes in many organisms, particularly within families *Planococcaceae* and *Erysipelotrichaceae.* Members of these families, such as *Lysinobacillus sphaericus* and *Clostridium innocuum*, could be excellent model organisms for studying sporulation mechanisms, such as engulfment, formation of the spore coat, and spore germination.

## INTRODUCTION

A variety of bacteria are capable of producing resting forms, commonly referred to as spores. However, the ability to form heat-, solvent- and UV-resistant endospores has only been observed in members of the phylum Firmicutes (low-G+C Gram-positive bacteria, recently renamed *Bacillota*) (1–8). Occasional reports of endospore-formers from other phyla have not been validated so far (9, 10). Sporulation enables bacteria to survive adverse environmental conditions. Thus, when this ability is found in human pathogens that cause severe diseases, such as anthrax, botulism, colitis, infectious diarrhea, gas gangrene, sepsis, tetanus, and food poisoning, it makes them particularly dangerous and difficult to eradicate (4, 5). Even spore-formers that used to be considered benign could turn out to be opportunistic pathogens, as exemplified by the recently described involvement of *Paenibacillus thiaminolyticus* in postinfectious hydrocephalus (11).

Formation of endospores is a complex process that starts with the asymmetric division of a vegetative cell producing a mother cell and a prespore, and proceeds through several stages of spore maturation. In the best-studied model organism *Bacillus subtilis*, the sporulation process affects expression of more than 500 genes, some of which are essential for sporulation, whereas others appear to be involved in various regulatory circuits (12–20). While some genes are exclusively involved in sporulation and are only expressed at certain stages of the process, spores also contain many housekeeping proteins that function during spore germination and subsequent vegetative growth (21–23). Studies on sporulation mechanisms in clostridia identified a somewhat smaller set of sporulation genes than in bacilli, leading to the conclusion that clostridia encode a streamlined version of the sporulation machinery (22, 24–27).

The ability to form spores is widespread in the two major classes of Firmicutes, *Bacilli* and *Clostridia*, and also has been observed in the other, more recently described, firmicute lineages (4, 28–30). However, some well-studied taxa within the *Bacilli*, such as lactobacilli, listeria, staphylococci, and streptococci, do not include any spore-forming members (4). Similarly, no spore-formers have been identified in the clostridial family *Halanaerobiaceae* (order *Halanaerobiales*) and in several families of the order *Clostridiales* (recently renamed *Eubacteriales*) (4, 31). Many other firmicute lineages include both spore-forming and non-spore-forming members (4).

These observations, coupled with the attempts to identify potential drug targets for extermination of spores, call for identification of the core set of essential sporulation genes that are necessary and, possibly, sufficient for a bacterium to be a spore-former. The most obvious candidate, the sporulation master regulator Spo0A, indeed, appeared to be essential, as no *spo0A*^−^ organism had been shown to form spores (4, 32). However, Spo0A is encoded in the genomes of many non-spore-formers and, therefore, could not be used as a reliable predictor of the sporulation ability (4, 32). Three more genes, *sspA, dpaA* (*spoVFA*), and *dpaB (spoVFB)*, initially proposed as sporulation signatures (33), presented the same problem, being present in a variety of non-spore-formers (32).

The availability of complete genome sequences of many diverse firmicutes offers an opportunity to identify sporulation genes through comparative genomics. Following the approach pioneered by Stragier (34), several studies took advantage of the constantly expanding genome list to define the conserved core of sporulation genes that are present in all (or at least most) known spore-formers (2, 32, 35–37). The resulting list included about 60 genes, many of which had been previously shown to be essential for sporulation because mutations in these genes caused sporulation arrest and/or production of immature spores. However, deletion of some other genes from the core set appeared to have only a minor effect on the sporulation efficiency, indicative of a substantial redundancy of the sporulation machinery. Indeed, in *B. subtilis*, many sporulation genes are found in two or more paralogous forms (37, 38).

Here, we used the latest update of the Clusters of Orthologous Genes (COG) database (39) to analyze the sporulation gene sets in the current genome collection of diverse firmicutes, including members of the recently defined classes *Negativicutes*, *Tissierellia*, *Erysipelotrichia*, and *Limnochordia*. The COG database covers a limited number of selected, completely sequenced microbial genomes (typically, a single representative per bacterial genus) and features COG-specific patterns of presence-absence of genes in the respective organisms (40, 41). Thus, COG profiles offer an easy way to identify those genomes in which a given gene, e.g. one involved in sporulation, is missing (39, 41, 42). Analysis of COG profiles of 180 genomes representing 76 spore-forming and 102 asporogenous species allowed us to detect numerous events of lineage-specific gene loss in various components of the sporulation machinery. Of particular interest was the widespread loss of sporulation genes in the families *Planococcaceae* and *Erysipelotrichaceae,* which resulted in a greatly streamlined machinery for engulfment, formation of the spore coat, and spore germination. These observations suggest that certain members of these families could be excellent model organisms for studying the fundamental mechanisms of sporulation in greater molecular detail.

## RESULTS

### Spore-former genome collection

The analyzed set consisted of 180 genomes from 160 genera, representing every named genus of Firmicutes that included at least one completely sequenced genome by April 1^st^, 2019, and several genomes released after that date (39). These organisms belong to six currently recognized classes of Firmicutes: *Bacilli, Clostridia, Erysipelotrichia, Limnochordia, Negativicutes*, and *Tissierellia*, and comprise 76 species whose original descriptions mentioned their ability to sporulate, 102 non-spore-formers (based on the descriptions of the respective strains), and two species with an unclear sporulation status (**Table S1** in the Supplemental material). At the time of this study, there were no completely sequenced genomes from any representatives of the classes *Culicoidibacteria* and *Thermolithobacteria*; members of these classes have been described as non-spore-forming (43, 44).

To ensure reliability of the gene presence-absence patterns, this work relied on high-quality genomes included in the recent release of the COG database (39), most of which had been vetted by the NCBI’s RefSeq team and selected as RefSeq reference genomes (45). For two organisms, *Caproiciproducens* sp. NJN-50 and *Thermincola potens* JR, the sporulation status had not been described and they had both spore-forming and asporogenous relatives. To ensure proper coverage of the class *Erysipelotrichia*, its two members featured in the COG database were supplemented with five additional genomes, including two from known spore-formers, [*Clostridium*] *innocuum* (the square brackets indicate that this organism has been misnamed, which remains to be rectified (28)) and *Erysipelatoclostridium* (formerly *Clostridium*) *ramosum* (28, 46, 47).

### Sporulation and genome size

As reported previously, spore-formers generally have larger genomes than non-spore-formers (32). In our set, the mean genome size of the former was ∼3.9 Mb, compared to ∼2.7 Mb for the latter (**Figure S1** in the Supplemental material). Although the genome size distributions overlapped, the previously reported 2.3-Mb boundary for cultivated spore-formers (32) held for the expanded genome set analyzed here. Only two spore-formers in the set, both still uncultured, *Candidatus* Arthromitus SFB-mouse and *Candidatus* Desulforudis audaxviator MP104C, had genome sizes of less than 2.4 Mb (see Table S1; the recently cultivated spore-forming strain of *Ca.* Desulforudis audaxviator with a 2.2-Mb genome (48) was not part of the COG set). In agreement with the previous reports (4, 49, 50), the ability to sporulate often differed even between closely related organisms: families *Bacillaceae* and *Planococcaceae* in the class *Bacilli*, most families in class *Clostridia*, and the families *Tissierellaceae* and *Erysipelotrichaceae* all included both spore-forming and asporogenous members (Table S1). A comparison of the genome sizes of spore-formers and non-spore-formers from different classes is presented in Figure S1B.

### Selection of sporulation genes

For the purposes of this work, the ‘sporulation genes’ were defined as those genes that participate in spore formation but have no known housekeeping roles in vegetative cells. This approach excluded most metabolic enzymes, as well as proteins involved in DNA replication and repair, transcription, translation, motility, secretion, and other processes. However, we included the cell division genes that are (also) involved in asymmetric cell division, as well as Spo0A-regulated genes that function at the onset of sporulation; pre-spore and forespore genes expressed under the control of SigF and/or SigG; mother cell genes that are expressed under the control of SigE and/or SigK; genes involved in the formation of the spore cortex and spore coat; and genes involved in spore germination. The list of the genes belonging to these groups was compiled based on the previous studies (2, 15, 16, 28, 32, 36, 37) and the data from the SubtiWiki database (http://subtiwiki.uni-goettingen.de/) (38). In the COG database, products of these genes formed 237 clusters of orthologs (COGs). We extracted COG phyletic profiles (the patterns of gene presence-absence in the respective genomes) for these sporulation genes and sorted them by their functions and representation among the 76 spore-forming organisms. The list of the most common sporulation genes identified in this manner is presented in **Table 1**. The patterns of distribution of these genes among all 180 analyzed species are listed in **Table S2** in the Supplemental material and are shown graphically in **Figure S2**.

**Table 1.**
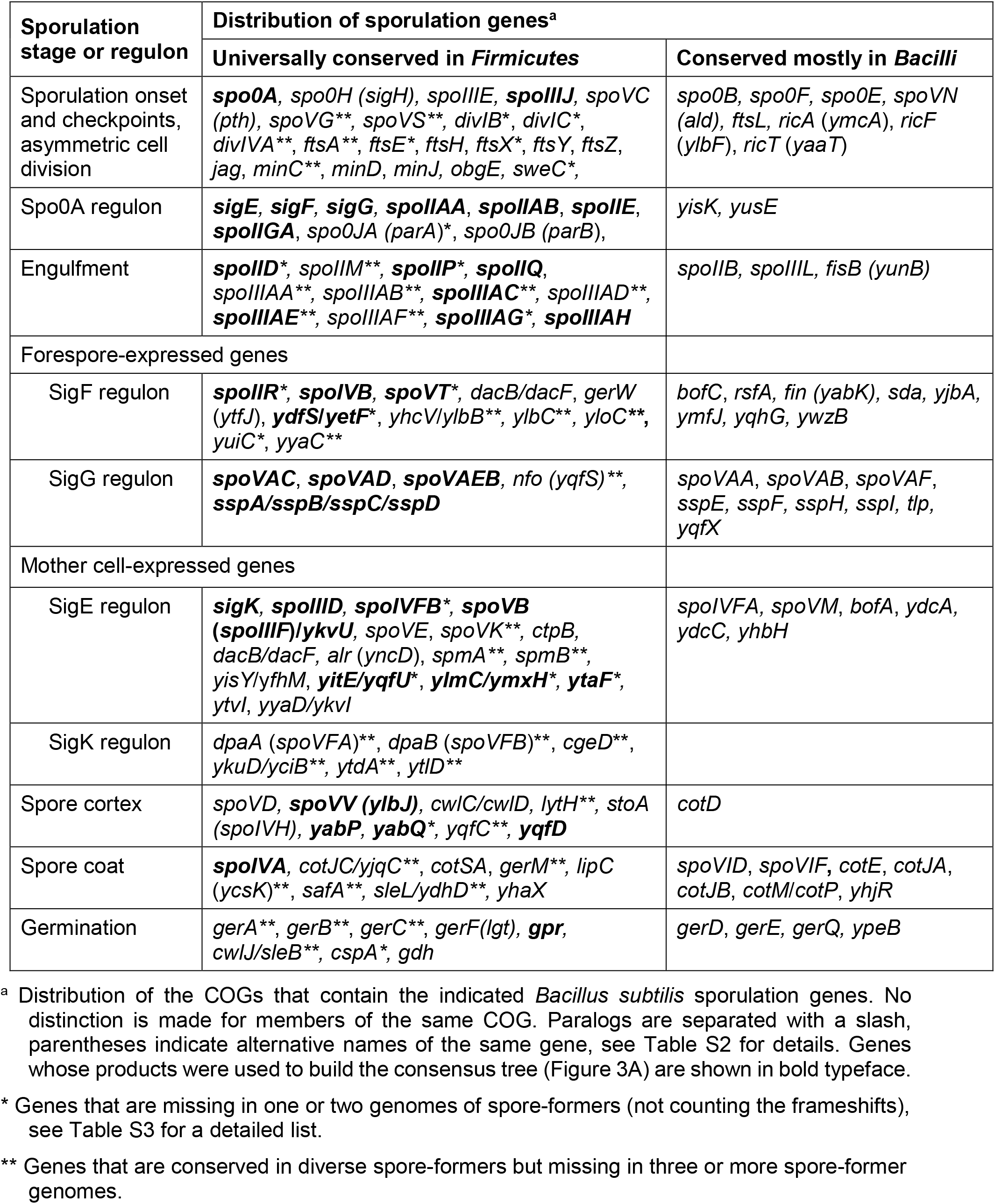
Conservation of sporulation genes in spore-forming Firmicutes.

### Distribution of *spo0A*, *dpaAB*, and *sspA* genes

In 2004, Weigel and colleagues identified four conserved sporulation genes, namely, *spo0A, dpaA, dpaB*, and *sspA*, and proposed using these genes as markers of the ability of a bacterium to form endospores (33). We have previously reported the presence of these genes in several clostridial non-spore-formers and argued that the presence of these genes was insufficient – and, in the case of *dpaAB*, unnecessary – for sporulation (32). It was instructive to test the distribution of these four genes in the current, much more diverse collection of firmicute genomes.

### spo0A

Analysis of the current genome set confirmed that the master cell regulator Spo0A is an essential component of the sporulation machinery. Indeed, the genome of every experimentally characterized spore-forming member of the Firmicutes encodes Spo0A, and its absence is an excellent predictor of the organism’s inability to sporulate. Among the 180 species analyzed in this work, the *spo0A* gene was present in 118, including all 76 spore-formers and 40 non-spore-formers (Table 1, Table S1). These observations confirm that the presence of *spo0A* is insufficient to conclude that a bacterium is a spore-former although its absence unequivocally indicates that it is not. In agreement with the previous reports (32, 33, 51), Spo0A-encoding non-spore-formers were found in classes *Bacilli*, *Clostridia*, and *Erysipelotrichia.* Among *Bacilli*, our set included six Spo0A-encoding non-spore-formers, all in the order *Bacillales*: *Lentibacillus amyloliquefaciens* in the family *Bacillaceae*, *Kurthia zopfii* and *Planococcus antarcticus* in *Planococcaceae*; *Macrococcus caseolyticus* in *Staphylococcaceae*, *Novibacillus thermophilus* in *Thermoactinomycetaceae,* and *Exiguobacterium* sp. from *Bacillales* Family XII Incertae Sedis. No Spo0A-encoding genomes and accordingly no spore-formers were represented among the 23 members of the order *Lactobacillales*. The current version of GenBank includes several *spo0A* genes from several clinical isolates of *Streptococcus pneumoniae*, but it remains unclear whether these isolates are correctly classified.

Spo0A is encoded by 32 of the 42 non-spore-forming members of *Clostridia* in our set and both members with an unclear sporulation status. Among the ten members of *Negativicutes* and nine members of *Tissierellia*, all four Spo0A-encoding organisms were spore-formers. Among the seven representatives of *Erysipelotrichia*, Spo0A was encoded in two spore-formers, *C. innocuum* and *E. ramosum*, and two non-spore-formers, *Amedibacterium intestinale* and *Turicibacter* sp. (Table S1). Most Spo0A^+^ genomes, 107 out of 118, carried a single *spo0A* gene. The only exceptions were observed in the order *Clostridiales* where 10 genomes encompassed two paralogous *spo0A* genes each, and one, the asporogen *Anaerostipes hadrus*, had three *spo0A* paralogs. Irrespective of their ability to form spores, Spo0A-encoding organisms had larger genomes than Spo0A^−^ ones (Figure S1B) and possessed many more sporulation genes than Spo0A^−^ genomes (**Figure 1**). This trend was most pronounced for widespread sporulation genes (Figure 1A) but also held for those sporulation genes that were conserved mostly in *Bacilli* (Figure 1B) and for more narrowly conserved sporulation genes (Figure 1C). The phylogenetic tree of Spo0A proteins (**Figure S3** in the Supplemental material) showed only minor deviations from the 16S rRNA-based phylogeny of the Firmicutes, suggesting vertical inheritance of the *spo0A* genes within this phylum (see below).

**Figure 1.**
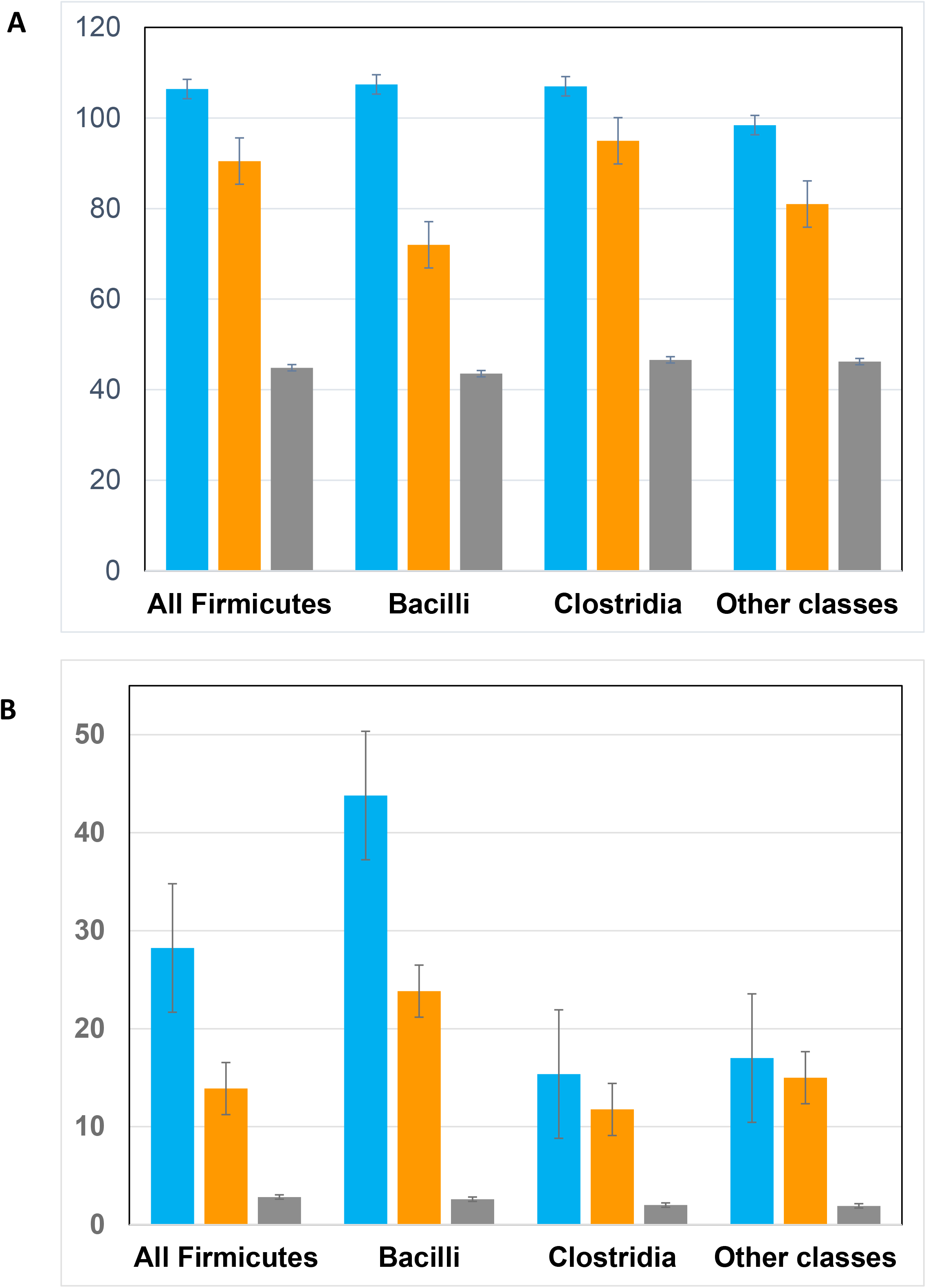
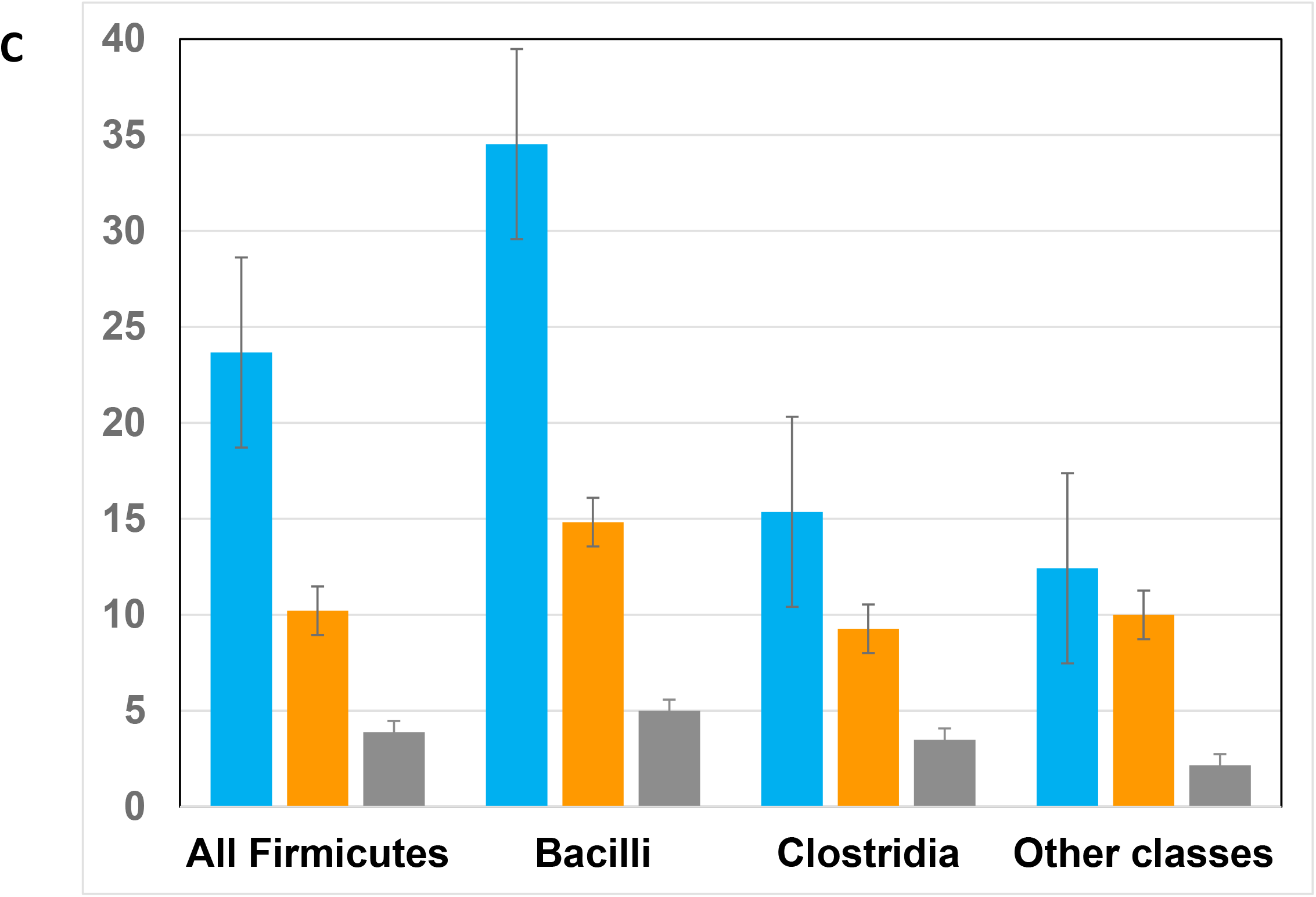
Distribution of sporulation genes in Spo0A^+^ and Spo0A^−^ firmicutes. The graphs show the numbers of widespread sporulation genes (A, out of 112), those conserved mostly in *Bacilli* (B, out of 51), and more narrowly conserved sporulation genes (C, out of 74) encoded in spore-formers (which are all Spo0A^+^, blue columns), Spo0A^+^ non-spore-formers (orange columns), and Spo0A^−^ non-spore-formers (gray columns). The column groups represent, from left to right, all firmicutes, bacilli, clostridia, and members of the other four classes (*Negativicutes*, *Tissierellia*, *Erysipelotrichia*, and *Limnochordia*).

### dpaAB

Dipicolinate, a key spore component, is produced by oxidation of dihydrodipicolinate, which is catalyzed by dipicolinate synthase, whose two subunits are encoded by *dpaA* (*spoVFA*) and *dpaB* (*spoVFB*) genes that are expressed in the mother cell under σ^K^ control. Among the members of the class *Bacilli* in our genome set, the *dpaAB* gene pair was found in all spore-formers and was absent in nearly all non-spore-formers. Therefore, within *Bacilli*, these genes indeed could serve as markers of sporulation. The *dpaAB* gene pair was also found in *Limnochorda pilosa* and in both spore-forming members of *Erysipelotrichia*. In clostridia, however, the picture was more complicated. The *dpaA* and *dpaB* genes have been previously shown to be missing in *Clostridium perfringens*, *C. botulinum*, and *C. tetani*, in which dihydrodipicolinate oxidation is catalyzed by the electron transfer flavoprotein, encoded by the *etfA-etfB* gene pair (32, 52). Non-orthologous displacement of *dpaAB* by *etfAB* was also observed in several other clostridial spore-formers, including *Clostridium acetobutylicum*, *Alkaliphilus metalliredigens*, *Paeniclostridium sordellii*, and *Tepidanaerobacter acetatoxydans*, as well as two spore-forming representatives of *Negativicutes*, *Methylomusa anaerophila* and *Pelosinus fermentans*, and in *Tissierella* sp. JN-28 (aka *Sporanaerobacter* sp. NJN-17), a member of the class *Tissierellia*. Finally, *Gottschalkia acidurici* does not encode either DpaAB or EtfAB, suggesting that in this organism dipicolinate production is catalyzed by yet another, currently unidentified oxidoreductase.

### sspA

The distribution of the *sspA* gene, which encodes α/β-type small acid-soluble sporulation protein (SASP), is also dramatically different between classes *Bacilli* and *Clostridia*. Among bacilli, *sspA* is found in all spore-formers and is missing in nearly all non-spore-formers (Table S2), making it another good marker of the ability of bacilli to sporulate. Essentially the same picture was observed in the classes *Negativicutes*, *Tissierellia*, and *Erysipelotrichia*. However, in *Clostridia*, the *sspA* gene is represented both in spore-formers and in many non-spore-formers, albeit only in Spo0A-encoding ones. The only spore-former in our set that did not carry the *sspA* gene was *C. innocuum*. The ability of this organism to form spores in the absence of SspA (or any other known SASPs; see below) merits further study.

### Identification of the core sporulation gene set

The COGs that included the *B. subtilis* sporulation genes were sorted by their representation in the 76 spore-former genomes and, specifically, in the genomes of spore-forming bacilli and clostridia. We classified these COGs into four groups (Table 1): i) genes (COGs) that were represented in every spore-former genome, with the possible exception of one or two; ii) genes that were widespread in spore-formers but were missing in three or more genomes, often those from a specific lineage; iii) genes that were widespread in bacilli but poorly represented in other classes, and iv) genes that were only present in a limited number of spore-formers. Representation of these genes (COGs) in all analyzed firmicute genomes was recorded as well (Table S2).

As mentioned above, the analyzed genomes were extracted from the COG collection (39), most of which were selected from the NCBI’s RefSeq database (45). Nevertheless, it has been shown that even highly conserved ribosomal proteins are occasionally missed during genome assembly, left untranslated because of sequencing errors, or simply overlooked in the course of genome annotation (53). Therefore, cases of widespread sporulation genes missing in one or more spore-former genomes were individually verified by checking for the potential presence of highly diverged or partial ORFs using DELTA-BLAST search (54) against the NCBI protein database and/or by TBLASTn search (55) against the nucleotide sequence database, each time selecting the respective species or strain as the target database and a representative protein from a closely related organism as a query (56). This analysis identified frameshifts and nonsense mutations in several widely conserved sporulation genes in several genomes (**Table S3** in the Supplemental material). However, the occurrence of such suspicious frameshifts was typically limited to one or two per genome, even in *Sulfobacillus acidophilus* str. TPY (GenBank accession number <u>CP003179.1</u>), which had several unannotated ribosomal proteins (53) and was excluded from RefSeq because of the missing rRNA genes. This genome also lacked an unusually high number of widespread sporulation genes (**Table S4** in the Supplemental material). Another exception was the genome of *Jeotgalibacillus malaysiensis* D5 (GenBank: CP009416.1), which also had several frameshifted and missing sporulation genes (Tables S3 and S4), some of which were found in other *Jeotgalibacillus spp.* (Table S2). The peculiar phylogenetic position of the genus *Jeotgalibacillus* between *Bacillaceae* and *Planococcaceae* (57, 58) warranted keeping this genome in the analyzed set but its sporulation genes patterns were considered unreliable.

Comparison of Table 1 with the previously identified sets of core sporulation genes (2, 32, 36, 37) showed that the list remained largely stable, despite the substantially expanded coverage of *Bacilli* and *Clostridia* and inclusion of representatives of three more classes of Firmicutes, *Negativicutes*, *Tissierellia*, and *Limnochordia* (**Table S5** in the Supplemental material). Most genes previously assigned to the conserved core were found in all or nearly all of the 76 spore-formers in the current genome set. Further, several genes, such as *spoIVFB*, *spoVE*, *cotS*, *yfhM*, *yhaX*, *yisY*, and *yyaA*, turned out to be more widespread than previously thought (Table S5). The only deviation from this pattern was observed in the class *Erysipelotrichia*, in which the spore-forming members, *C. innocuum* and *E. ramosum*, missed more than a dozen genes from the conserved sporulation gene set (see below). This comparison also showed that, despite the overall conservation of the core sporulation genes in all classes of Firmicutes, many of these genes were found to be completely missing (as opposed to being disrupted by frameshifted) in one, two, or more genomes (**Table S6** in the Supplemental material). Although some of these gaps in phyletic patterns could be caused by errors in genome sequencing and/or assembly, the consistency of such patterns in certain phylogenetic groups appeared to reflect an actual evolutionary trend. In the following sections, we describe the streamlining of the sporulation gene complement in spore-formers from families *Planococcaceae* and *Erysipelotrichaceae*, followed by the discussion of the gene loss within specific sporulation systems.

### Sporulation genes in *Planococcaceae*

The analyzed genome set included six spore-forming members of the family Planococcaceae (also known as Caryophanaceae), order Bacillales: Paenisporosarcina sp. K2R23-3, Rummeliibacillus stabekisii, Solibacillus silvestris, Sporosarcina psychrophila, and Ureibacillus thermosphaericus. In addition, we included Jeotgalibacillus malaysiensis, which is currently assigned to the Planococcaceae, and Lysinibacillus sphaericus, which is usually assigned to Bacillaceae but in phylogenetic analyses robustly falls within the Planococcaceae (58). These seven organisms showed a consistent pattern of sporulation gene loss, with most of them lacking four of the nine components of the SpoIIQ–SpoIIIA complex (spoIIIAA, spoIIIAB, spoIIIAD, and spoIIIAF genes), as well as such conserved sporulation genes as spmA, spmB, spoIIM, gerM, gerW (ytfJ), yqfC, and ytxC, which are found in most spore-formers (Tables 1, S2, S4). Furthermore, these seven organisms also lacked some genes that are generally conserved among the spore-forming members of the Bacillaceae: bofC, csfB, spoIIIL, spoIVFA, spoVAA, spoVAB, spoVAEA, spoVID, yabG, yhbB, yrrD, ysxE, and ytxC (Table S2). They also lack the genes that encode such SASPs as SspK, SspL, SspN, SspO, SspP, and CsgA; spore coat proteins CotM, CotI, CotO, CotP, and CotW, and spore maturation proteins SpsJ and CgeB, which are found in most spore-forming bacilli (Table S2). The most dramatic deviations from the common phyletic pattern were detected in J. malaysiensis, in accordance with its uncertain phylogenetic position (58). The consistent absence of all these genes suggests that members of the Planococcaceae possess substantially streamlined sporulation machinery. The analyzed genome set also included two asporogenous representatives of Planococcaceae, K. zopfii and P. antarcticus, which have spo0A but lack almost all other sporulation genes (Table S2).

### Sporulation genes in *Erysipelotrichaceae*

The evolution of the *Erysipelotrichia* lineage likely involved massive loss of genes, including many related to sporulation (59). Indeed, the genomes of *C. innocuum* and *E. ramosum*, despite their relatively large size (4.7 and 3.25 Mb, respectively), show patterns of extensive loss of sporulation genes. Consistent with the previous observations (59, 60), they both lack seven of the nine genes of the SpoIIQ–SpoIIIA complex, retaining only SpoIIQ and SpoIIIAH components. Further, they both lack such widely conserved sporulation genes as *spoIIID*, *bofA*, *cotQ*, *safA*, *yabQ*, *ypeB*, *yyaC*, *yyaD*, *sleL* (*yyaH*), and *gerW* (*ytfJ*) (Table S2, Table S4). Despite the evolutionary proximity of *Erysipelotrichia* and *Bacilli* (28, 59), many widely conserved bacillar sporulation genes are missing as well, including *bofC*, *csfB*, *spoIIIL*, *spoIVFA*, *spoVAA, spoVAB, spoVAEA*, *spoVID*, *yabG*, *yhbB*, *yrrD*, *ysxE, and ytxC*. Some sporulation genes appear to have been lost in only one of these genomes. In particular, *C. innocuum* lost *spoIIR* and *spoVS*, whereas *E. ramosum* retained these genes but lost *spoVG* and *spoVT* (Table S4). Another system with an unusual phyletic pattern is the spore germination receptor GerABC. While non-spore-forming members of *Erysipelotrichia* either encode all three subunits of GerABC (*Turicibacter* sp. H121) or none at all (three other genomes), the spore-formers *C. innocuum* and *E. ramosum* encode the membrane subunit GerA (GerAA) but not the membrane transporter GerB (GerAB) or the lipoprotein GerC (GerAC) subunits. Although the GerABC receptors are not strictly required for germination (they are missing in *Clostridioides difficile* (3, 61) and could be dispensable in *C. botulinum* (62)), they are nearly ubiquitous among spore-formers (Table S2). The presence of GerAA but not GerAB or GerAC subunits is so far unique among known spore-formers.

### Sporulation onset and asymmetric division

Along with *spo0A*, genes involved in sporulation onset and asymmetric division are highly conserved in spore-formers of the class *Bacilli*, and many, although not all, of these genes, are also conserved in spore-forming members of *Clostridia* and four other classes of *Firmicutes* (Table 1, Table S2). As discussed previously, many clostridia lack the Spo0B-Spo0F-Spo0A phosphorelay system, and their Spo0As are phosphorylated by orphan histidine kinases that are not orthologous to any of the five sporulation histidine kinases (KinA – KinE) of *B. subtilis*. However, many of these orphan kinases contain at least one PAS sensor domain (25, 51, 63). Nevertheless, the *spo0B*-*spo0F* gene pair is present in several clostridia, primarily members of the families *Heliobacteriaceae*, *Peptococcaceae*, and *Thermoanaerobacteraceae* (51), as well as in *Negativicutes* and *Tissierellia* (Table S2).

The genes involved in asymmetric cell division, such as *divIB, divIC, divIVA, ftsE, ftsH, ftsX, ftsY, ftsZ, minC, minD, and minJ*, are widespread in bacteria, including most firmicutes and nearly all spore-formers (Table S2). As expected, genes encoding the four sporulation-specific sigma subunits, *sigF*, *sigG*, *sigE* and *sigK*, are found in every spore-former, but the Spo0A-regulated genes *spoIIAA*, *spoIIAB*, *spoIIE*, *spoIIGA*, *spo0JA* (*parA*), and *spo0JB* (*parB*) are also highly conserved in spore-formers of every firmicute lineage (Table 1). However, such Spo0A-dependent genes of *B. subtilis* as *ykuJ*, *ykuK*, and *yneF*, all three with unknown functions, are not conserved even within *Bacillales* (Table S2).

### Formation and activation of σ^K^

While σ^K^ is universally encoded in the genomes of all spore-forming firmicutes, its activation is tightly controlled, preventing premature expression of late-stage sporulation genes in the mother cell (64–66). In *B. subtilis*strain 168, the σ^K^-encoding gene is interrupted by a 48-kb prophage-like *skin* (*<u>s</u>ig<u>K</u>*<u>-in</u>tervening) element, which results in two separate coding regions denoted *spoIVCB* and *spoIIIC*, neither of which is expressed. These fragments of *sigK* are joined as a result of SpoIVCA-dependent excision of *skin* to form the complete, intact *sigK* gene that encodes pro-σ^K^ (67, 68). Pro-σ^K^ is then activated in the mother cell through cleavage by the intramembrane metalloprotease SpoIVFB (64, 66). In *C. difficile* 630, the *skin* element is only 14.6 kb long, and its excision yields a *sigK* gene encoding a mature σ^K^, which does not require proteolytic activation (69, 70). SigK-interrupting *skin* elements have also been identified in *C. tetani* and *C. perfringens* (71, 72). We confirmed the presence of split *sigK* genes in *B. subtilis*, *C. difficile,* and *C. tetani,* and identified split *sigK* genes in 12 additional organisms from *Bacilli*, *Clostridia*, *Negativicutes*, and *Tissierellia* (**Figure 2**). These *skin* elements range in size from 2.6 to 48 kb and encode from 2 to 67 proteins (see **Table S7** in the Supplemental materials).

**Figure 2.**
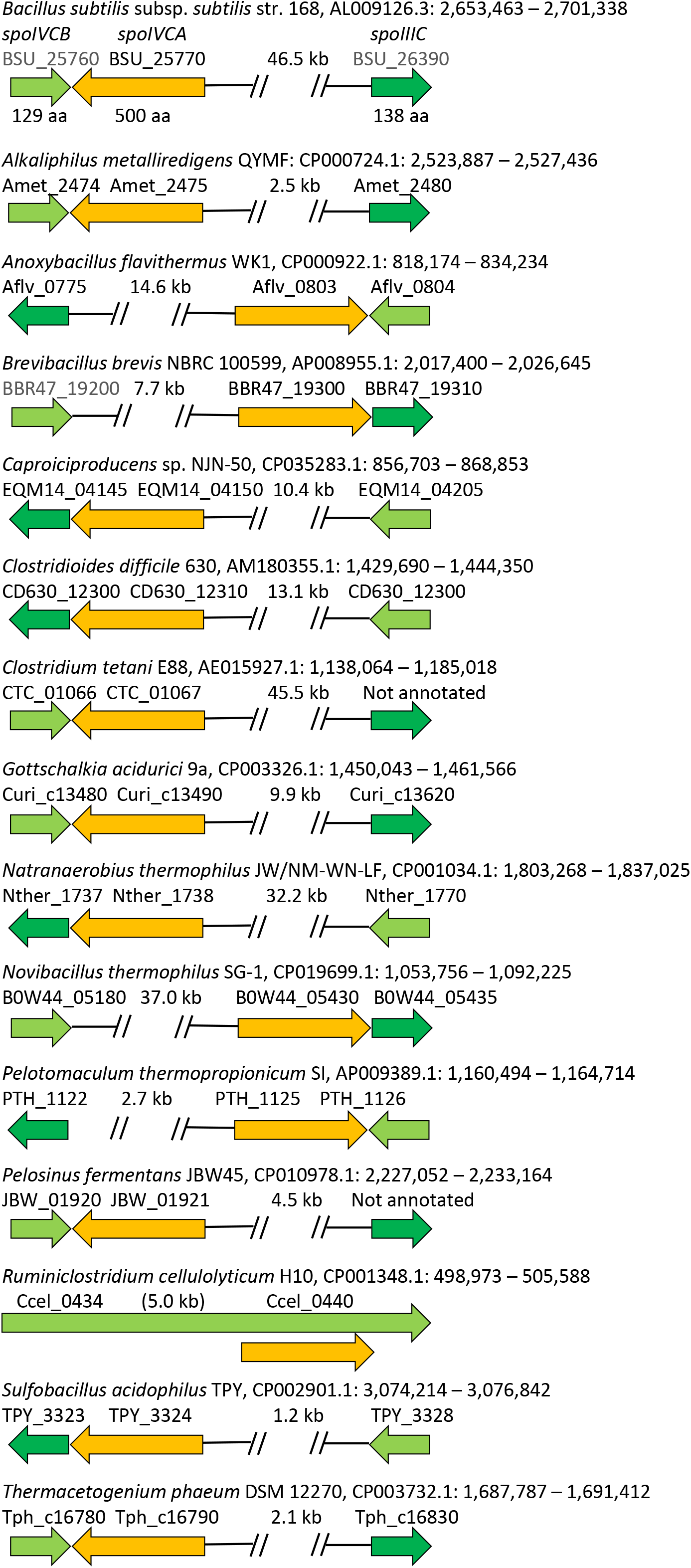
Organization of *skin* elements in 15 firmicute genomes. *spoIVCB* genes are shown in light green, *spoIIIC* in dark green, *spoIVCA* in orange. The organism names are followed by the GenBank genome entries and genomic coordinates of the regions between *spoIVCB* and *spoIIIC.* The numbers above the gaps indicate the distances between *spoIVCB* or *spoIIIC* and *spoIVCA* (∼1.5 kb less than the total length of the *skin* element). *R. cellulolyticum* Ccel_0434 is annotated as a single 7,394-nt pseudogene.

The orientation of skin elements with respect to the *sigK* component genes is variable, but the *spoIVCA* gene is always located at the edge of the element, usually immediately upstream of the *spoIIIC* gene, which encodes the C-terminal fragment of σ^K^ (Fig. 2). The SpoIVCA proteins from different *skin* elements (see **Figure S4A** in the Supplemental materials for a sequence alignment) are members of a single family of site-specific DNA recombinases (COG1961) that is represented by multiple copies in most firmicute genomes, including up to 26 genes in (non-spore-forming) *Flavonifractor plautii* and *Oscillibacter valericigenes.* In the phylogenetic tree of SpoIVCA-like recombinases/integrases (**Figure S4B**), SpoIVCAs from *skin* elements are interspersed with those from tailed phages (Caudovirales), suggesting (pro)phage origin of the *skin* elements. Remarkably, aside from *spoIIIC, spoIVCA* and *spoIVCB,* no genes are shared by all *skin* elements (**Figure S4C**). The presence of *skin* elements that split *sigK* genes in two non-adjacent fragments often causes confusion in genome annotation. Thus, in the current genomic entries for *B. subtilis* strain 168 (Genbank accessions AL009126.3, CP051860.2, CP053102.1, and CP052842.1), *sigK* is erroneously marked as a pseudogene, whereas in genomic entries for *C. tetani* E88 (GenBank: AE015927.1) and *Pelosinus fermentans* (GenBank: CP010978.1), the C-terminal *spoIIIC*-like fragment is left untranslated (Tables S5, S7). Remarkably, *sigK* is not unique as the *skin* insertion site: *skin*-like elements were also found interrupting the *cotJC* gene of *Laceyella sacchari* and *spoIIID* gene of *Ruminiclostridium cellulolyticum* (Table S3).

In *B. subtilis,* activation of pro-σ^K^ by SpoIVFB is regulated by BofA, BofC, and also SpoIVFA, which is itself regulated by proteolysis by the forespore-expressed protease SpoIVB (73). σ^K^ is encoded in every spore-former genome as is, with a single exception, SpoIVFB, which is a member of a widespread metalloprotease family (COG1994). However, SpoIVFB-interacting BofA, BofC, and SpoIVFA are far less widespread. BofA is missing in 10 of the 76 spore-formers, including six clostridia, *C. innocuum*, *E. ramosum*, and *Limnochorda pilosa.* The distribution of BofC and SpoIVFA is generally limited to the members of the class *Bacilli*. The full-length *B. subtilis*-like BofC, which consists of two distinct domains (74), is found almost exclusively in the families *Bacillaceae* and *Paenibacillaceae*; it is missing in some members of the former and, conspicuously, in members of the *Planococcaceae* (Table S2). The C-terminal domain of BofC, PF08955, is more widespread but is still missing in most spore-forming clostridia. SpoIVFA is missing in the *Planococcaceae*, in nearly all clostridia, and, again, in *C. innocuum*, *E. ramosum*, and *L. pilosa.* Remarkably, SpoIVFA-cleaving serine protease SpoIVB is nearly universal among spore-formers, being represented in numerous genomes that do not encode SpoIVFA. The details of SpoIVFB–BofA interaction in the absence of SpoIVFA, and the functions of SpoIVB homologs in these organisms remain to be elucidated.

### Conservation of the engulfment complex

In *B. subtilis*, engulfment is triggered by SpoIIB and involves two key protein complexes, SpoIID-SpoIIM-SpoIIP (the DMP complex) and SpoIIIA-SpoIIQ, which are targeted to the growing septum by either SpoIIB or the SpoIVFA-SpoIVFB pair (75–77), with additional involvement of GerM (78). Both protein complexes are found in *C. difficile* (79) and are conserved in almost all spore-formers (Table 1), whereas the targeting proteins are less widespread: as noted previously (32), both *spoIIB* and *spoIVFA* are missing in clostridia (Table S2).

Among the three genes encoding the peptidoglycan-degrading DMP complex, *spoIID* was missing in a single genome in our 76 spore-former genome set, and *spoIIP* was missing in two (Table S6); these three cases could be due to the sequencing problems. In contrast, *spoIIM*, although widespread as well, was missing in nine genomes (Table S2). This pattern is consistent with the reports that SpoIIM is not essential for engulfment in *C. difficile* (80, 81), whereas SpoIID and SpoIIP appear to be indispensable.

The transmembrane SpoIIIA–SpoIIQ complex consists of nine proteins, eight of which are encoded in the mother cell by the genes of the *spoIIIAA-AB-AC-AD-AE-AF-AG-AH* operon that is under the SigE control. The ninth protein, SpoIIQ, is produced in the forespore under SigF control and contains the M23-type metallopeptidase LytM domain (82, 83). The core of the SpoIIIA-SpoIIQ complex consists of SpoIIIAH and SpoIIQ proteins. Both these proteins form dodecameric (or even larger) rings that interact to form a 60 Å (or even a 140 Å) transmembrane channel, which connects the mother cell and the forespore and enables trafficking of small molecules between the two compartments (84, 85). The structure of this complex from *B. subtilis* (**Figure S5A**) revealed a key role of the LytM domain of SpoIIQ in its interaction with SpoIIIAH. However, despite the presence of the common LytM domain, SpoIIQ proteins from bacilli and clostridia are substantially distinct: the M23 peptidase is inactivated in the former but appears to be intact in the latter (86). Accordingly, these proteins belong to two different COGs, COG5820 and COG5821, in the COG database (39). Both varieties of SpoIIQ are often encoded in similar *spoIID–spoIIQ– spoIIID* operons although the genomic neighborhoods of *spoIIQ* are highly variable even within the *Bacillaceae* (**Figure S6**). Although we previously reported not being able to find orthologs of clostridial *spoIIQ* (*CD0125* in *C. difficile)* in the genomes of several members of the family *Peptococcaceae* (32), in this work, using synteny and DELTA-BLAST (54) searches, we identified these in the genomes of all clostridial spore-formers but very few non-spore-formers (Table S2). Spore-forming members of *Erysipelotrichia*, *C. innocuum* and *E. ramosum*, encode the bacillar form of SpoIIQ, whereas spore-forming members of *Negativicutes*, *Tissierellia*, and *Limnochordia* encode the clostridial form. Thus, all 76 spore-formers in our set encode both SpoIIIAH and one of the two forms of SpoIIQ, indicating strict conservation of the SpoIIQ–SpoIIIAH pore among spore-forming bacteria.

As noted above, two groups of Firmicutes, *Erysipelotrichaceae* and *Planococcaceae*, have apparently lost multiple components of the SpoIIQ–SpoIIIA complex. The *Erysipelotrichia* members *C. innocuum* and *E. ramosum* retain only the *spoIIIAH* and *spoIIQ* genes. In addition, they possess SpoIVFB and GerM, but lack SpoIIB and SpoIVFA (Table S2, **Figure S5B, C**), see also reference (60). Six members of the family *Planococcaceae* show stronger conservation of the SpoIIQ–SpoIIIA complex: in addition to SpoIIIAH and SpoIIQ, they retain SpoIIIAC, SpoIIIAE, and SpoIIIAG components (Fig. S3). However, all these organisms lack SpoIIB, SpoIVFA, SpoIVFB, and GerM proteins that participate in the assembly of the SpoIIQ–SpoIIIA complex in *B. subtilis*. Members of *Planococcaceae*, such as *Lysinibacillus sphaericus*, could be useful model organisms for studying the core mechanisms of engulfment in the Firmicutes.

### Small acid-soluble sporulation proteins (SASPs)

SASPs are short proteins that bind double-stranded DNA and protect spores from heat, UV radiation, and other adverse conditions (87–91). In *B. subtilis*, the majority of the SASPs come from two families, α/β (COG5852, which unifies SspA, SspB, SspC, and SspD families, and COG5854, SspF) and gamma (COG5853), whereas several minor SASPs (87, 90, 92) comprise COGs from COG5855 through COG5864. As observed previously, the α/β-type SASPs are nearly universal in spore-forming firmicutes, whereas the distribution of other SASP families varies (32, 93). This observation is supported by the data from our new set. Indeed, COG5852 is represented in 75 of the 76 spore-formers and in 31 of the 40 Spo0A-encoding non-spore-formers (Table S2). The only spore-former missing SspA was *C. innocuum*; the other spore-forming member of *Erysipelotrichaceae*, *E. ramosum*, has two *sspA*-like paralogous genes and a single *sspI* gene for a minor SASP. Homologs of *sspA* are also found in other members of *Erysipelotrichaceae*, such as [*Clostridium*] *spiroforme* and [*Clostridium*] *cocleatum* (**Figure S7**). However, *C. innocuum* lacks known *ssp* genes, so the nature of its SASPs, if any, remains enigmatic. Most of the minor SASPs described in *B. subtilis* are also encoded in at least some members of *Bacillales* (93). In contrast, members of other classes of Firmicutes only encode α/β-type SASPs of the SspF and Ssp4 families (94, 95); some clostridia also encode Tlp-like SASPs. Thus, the diversity of SASPs in *B. subtilis* does not appear to be shared by other groups of Firmicutes, which is consistent with the reports of non-essentiality of minor SASPs (88, 95).

### Spore cortex

Genes involved in the biosynthesis of the spore cortex, the peptidoglycan layer that surrounds the forespore membrane, have been previously found to be conserved throughout *Bacilli* and *Clostridia* (32). Examination of the current expanded genome set showed that this conclusion still holds for *spoVB*, *spoVD (ftsI)*, *spoVE (ftsW)*, *spoVV* (*ylbJ)*, *yabP*, and *yqfD* genes (Table 1). One more gene, *yabQ*, which is essential in *B. subtilis* (96, 97), was only missing in the genomes of *C. innocuum* and *E. ramosum,* whereas the *yqfC* gene was missing in the genomes of six *Planococcaceae* members and *E. ramosum* (**Table S6**).

### Spore coat

As noted previously (32, 98, 99), bacilli and clostridia have dramatically different spore coats: many of the ∼70 proteins that form the coat of *B. subtilis* (7) are only conserved within *Bacillaceae*, or not even in all members of this family (32, 100). Among the ten morphogenetic spore coat proteins of *B. subtilis* (SpoIVA, SpoVM, SpoVID, SafA, CotE, CotH, CotO, CotX, CotY, and CotZ), only SpoIVA is universally conserved in all spore-forming firmicutes (Table S2). In bacilli, SpoIVA interacts with SpoVID (101, 102), whereas in *C. difficile* and some other clostridia, its interaction partner is SipL (CD3567), which shares with SpoVID a common C-terminal LysM domain (103, 104). Of the other SpoIVA- and/or SpoVID-interacting proteins, SpoVM is found in some clostridia but is not essential for sporulation (105), CotE is found only in a few clostridia, and the full-length SafA (as opposed to its LytM domain) is not represented outside of bacilli. CotH, CotO, CotX, CotY, and CotZ are missing even in some members of *Bacillaceae* and, except for CotH, none of these genes is found in clostridia or members of other classes of *Firmicutes* (Table S2).

Among the proteins of *B. subtilis* inner coat, the most widespread are proteins of the CotJC/YjqC family, which belong to the COG3546 “Mn-containing catalase”. Three members of this family (CotC, CotD, and CotE) were found in the spore coat of *C. difficile*, and the latter two have been shown to retain enzymatic activity (98, 106). Two more spore coat proteins of *C. difficile*, peroxiredoxin CotE and superoxide dismutase SodA, could be involved in protection from oxidative stress (98). Other widely conserved proteins of the *B. subtilis* basement layer and inner coat are also (not necessarily active) enzymes: acetyltransferase CgeE, peptidoglycan hydrolases CwlJ and SleL (YaaH), phospholipase LipC, HAD family phosphatase YhaX, ATP-grasp family enzymes YheC and YheD, MenH-related esterase YisY, and glycosyltransferases CotSA and YdhD (Table S2). A similar pattern is apparent in the outer coat, which contains alanine racemase YncD, nucleotidyl transferase YtdA, predicted FAD-dependent dehydrogenase CotQ, and LysM domain-containing YkzQ. Both inner and outer coats of *B. subtilis* contain members of the small heat shock protein (HSP20) family, CotP and CotM, respectively. These examples demonstrate the widespread recruitment (exaptation) of common housekeeping enzymes as structural components of the spore coat. Their broad representation among bacillar spore-formers, as opposed to the scarcity in non-spore formers (Table S2), suggests that such recruitment is a common mechanism of spore coat assembly in *Bacilli*. There is still insufficient data on spore coat proteins in *Clostridia* and none on those in other classes of *Firmicutes,* but studies in *C. difficile* (98, 106) suggest that such recruitment occurs in clostridia as well.

### Germination proteins

As mentioned above, the common germination receptors consisting of GerA, GerB, and GerC components are encoded in all classes of Firmicutes (Table S2). In our set of 76 spore-formers, the only exceptions were two members of the family *Peptostreptococcaceae*, *C. difficile* and *Paeniclostridium sordellii*, which lacked all three components of the receptor, and two members of *Erysipelotrichaceae*, *C. innocuum* and *E. ramosum*, which lacked GerB and GerC. In *C. difficile*, the bile acid germinant receptor has been identified as the pseudoprotease CspC (99, 107, 108), a member of the vast family of subtilisin-like serine proteases (COG1404). The presence of multiple members of this protein family in most spore-forming firmicutes suggests that the CspC-dependent germination mechanism is not limited to *C. difficile* and could potentially function in other organisms, as an addition to the better-studied GerABC-mediated signaling.

The germination process involves degradation of the spore protective layers, the cortex, and the inner and outer coats, as well as hydrolysis of DNA-shielding SASPs. In *B. subtilis*, hydrolysis of the spore cortex is catalyzed by peptidoglycan hydrolases CwlJ and SleB, members of the same COG3773, which is widespread in spore-forming bacilli and clostridia and is also found in spore-forming members of *Tissierellia* and *Limonochordia*, albeit not in the *Negativicutes* or *C. innocuum* (Table S2). The regulators of the activity of these enzymes, however, are less widespread. YpeB, which regulates SleB activity in bacilli and many clostridia (109–111), is missing in *C. difficile* and several other clostridia, as well as in all members of *Negativicutes* and *Erysipelotrichia* in the current set (Table S2). In *C. difficile*, the principal spore cortex-hydrolyzing enzyme is SleC (CD630_05510), a multidomain protein that consists of a structurally, but not functionally, characterized DUF3869 domain [Pfam (112) domain PF12985] (113), a SpoIID-like amidase domain, and a C-terminal peptidoglycan-binding (PF01471) domain. SleC homologs are found in many clostridia, as well as *C. innocuum* and *E. ramosum*.

Degradation of SASPs is catalyzed by the germination protease, Gpr, an aspartate protease (95, 114) that is universally conserved in spore-forming firmicutes (Table 1) but is absent in non-spore-forming members of *Bacilli*, *Negativicutes*, *Tissierellia*, and *Erysipelotrichia*. Among the clostridial non-spore-formers, the distribution of Gpr is generally similar to that of Spo0A: Spo0A-encoding spore-formers usually have the *gpr* gene, whereas those genomes that lack *spo0A* also lack *gpr* (Table S2).

Among other genes involved in spore gemination in *B. subtilis*, only a few, such as *spoVAD*, *gdh*, *gerF* (*lgt*), *ykvU* and *ypeB*, are widespread, whereas others, such as *csgA*, *gerD, gerE, gerPA/gerPF, gerPB, gerPC, gerPD, gerPE*, and *spoVAEA*, are found mostly (or exclusively) in spore-forming bacilli (Table S2).

### Uncharacterized sporulation proteins

Despite numerous studies on sporulation in *B. subtilis* and other model organisms, functions of several widespread sporulation genes have been clarified only recently whereas functions of several others remain unknown (Table 1). Altogether, the SubtiWiki database (38) lists >250 *y*-genes that appear to be specifically expressed during sporulation (15, 16, 20, 115, 116), but have not been characterized experimentally. Assignment of such genes to the COGs allowed making at least tentative functional predictions for several *y*-genes (**Table S8** in the Supplemental material). Examples include YdgB and YteA proteins, both annotated as ‘uncharacterized” in UniProt, which belong to the COG1734 “RNA polymerase-binding transcription factor DksA“, and the uncharacterized proteins YqcK and YqjC that fall into COG0346 “Catechol 2,3-dioxygenase-related enzyme, vicinal oxygen chelate (VOC) family”. Experimental characterization of these genes and other widespread genes of unknown function could provide further insight into the details of sporulation mechanisms in diverse firmicutes.

### Evolution of sporulation

We employed the updated list of core sporulation genes (Table 1) to investigate the evolution of sporulation and compare it with the overall phylogeny of Firmicutes. We started by constructing a phylogenetic tree for Spo0A, the master regulator of sporulation that is encoded in each of the 76 spore-formers in our genome collection, as well as 40 non-spore-formers and 2 organisms with uncertain sporulation status (Table S1). Several attempts to construct maximum-likelihood trees from an alignment of full-length Spo0A sequences using MEGA X (117), with several different sets of parameters, resulted in trees where proteins from the members of the same bacterial family typically formed separate clades. However, these trees could not be considered fully reliable because many branches were weakly supported (data not shown). By contrast, removing the variable-length linkers connecting the N-terminal receiver (REC) domain of Spo0A with its DNA-binding domain and using IQ-TREE 2 (118), which automatically selected the best-fit model (119), resulted in a well-supported tree (Figure S3). Again, Spo0As from the members of the same bacterial family typically formed separate clades. Among the *Bacilli*, monophyly was observed for the families *Alicyclobacillaceae* and *Planococcaceae*, Spo0As from members of *Bacillaceae* and *Paenibacillaceae* each formed two separate clusters (Figure S3). In *Clostridia*, Spo0As from most members of the families *Clostridiaceae*, *Lachnoospiraceae* (formerly known as *Ruminococcaceae* or *Hungateiclostridiaceae*), and *Peptostreptococcaceae* (*C. difficile* and *Paeniclostridium sordellii*), as well as from the order *Halanaerobiales*, appeared to be monophyletic. Spo0As from *Peptococcaceae* and *Oscillospiraceae* each formed two separate clusters, while Spo0As from *Thermoanaerobacteraceae* were found through the entire clostridial branch. At the class level, Spo0As from spore-forming members of the *Erysipelotrichia* formed a strongly supported clade that mapped within the *Bacilli*, while members of the *Negativicutes* (family *Sporomusaceae*) and *Tissierellia* mapped within the clostridial clade (Figure S3).

Notably, the monophyly of Spo0As from the members of the same family did not depend on whether they came from spore-formers or non-spore-formers. Thus, Spo0As from all members of *Planococcaceae* formed a single clade, with Spo0As from non-spore-formers *Kurthia zopfii* and *Planococcus antarcticus* grouping together with Spo0As from planococcal spore-formers (Figure S3). Likewise, Spo0A from the spore-former *Lachnoclostridium phytofermentans* belongs to a clade with non-spore-forming members of the family *Lachnospiraceae*, whereas Spo0A from the non-spore-former *Dehalobacter* sp. CF mapped together with spore-forming members of the family *Peptococcaceae* (Figure S3).

To gain further insight into the evolution of sporulation, and Firmicutes in general, we constructed a ribosomal proteins-based phylogenetic tree for all 180 analyzed organisms as well as sporulation proteins-based and ribosomal proteins-based trees for the 76 spore-formers. The maximum-likelihood phylogenetic tree of 180 firmicutes (**Figure S8**) was built using IQ-TREE 2 (118) from a concatenated alignment of 54 ribosomal proteins (6951 total positions, 6366 unique) essentially as described previously (28, 120). This tree had excellent bootstrap support and largely agreed with the currently accepted 16S rRNA gene-based phylogeny of the Firmicutes. Members of most previously defined Firmicute families clustered together, typically as clades. The notable deviations from the current taxonomy (56) included members of the class *Erysipelotrichia* (family *Erysipelotrichaceae*), mapped within *Bacilli* as a sister group to *Listeriaceae* and *Lactobacillales,* and members of three other classes mapping within *Clostridia* as follows: a) all three families of class *Tissierellia* as a sister group to *Peptostreptococcaceae*; b) members of the class *Negativicutes* as a sister group to families *Peptococcaceae* and *Syntrophomonadaceae*, and c) *Limnochorda pilosa*, the sole member of the class *Limnochordia,* as a sister group to *Sulfobacillus acidophilus* and *Thermaerobacter marianensis* (*Eubacteriales* Family XVII. Incertae Sedis). This tree also suggested placement of some organisms with previously obscure taxonomy: *Intestinimonas butyriciproducens* mapped into *Oscillospiraceae*, whereas *Ndongobacter massiliensis* and *Ezakiella massiliensis* fell within the radiation of the family *Peptoniphilaceae* (class *Tissierellia*).

For a bird’s eye view of the phylogeny of sporulation, we constructed a maximum-likelihood tree based on a concatenated alignment of 41 core sporulation proteins (as listed in Table 1) from all 76 spore-formers in the current set (**Figure 3A**) and compared it with the ribosomal proteins-based tree for the same 76 species (**Figure 3B**). This comparison showed a strong, clear trend: with a few exceptions, sporulation protein sets from members of the same family formed well-supported clades. As an example, members of the *Planococcaceae* in both trees formed a sister group to the *Bacillaceae*, the only differences being the branching order of *Sporolactobacillus terrae*, a member of the family *Sporolactobacillaceae*, and *Alkalihalobacillus* (formerly *Bacillus*) *halodurans*, and two members of the *Erysipelotrichaceae*, *C. innocuum* and *E. ramosum*. The clostridial portions of both trees had closely similar topologies with essentially the same branching order for most groups. Members of the recently recognized classes *Negativicutes* (family *Sporomusaceae*), *Tissierellia*, and *Limnochordia* mapped within the clostridial lineage, but, again, in the same positions on both trees (Figure 3). The congruence of both trees indicated that the sporulation genes largely evolved by vertical inheritance, without substantial horizontal gene transfer.

**Figure 3.**
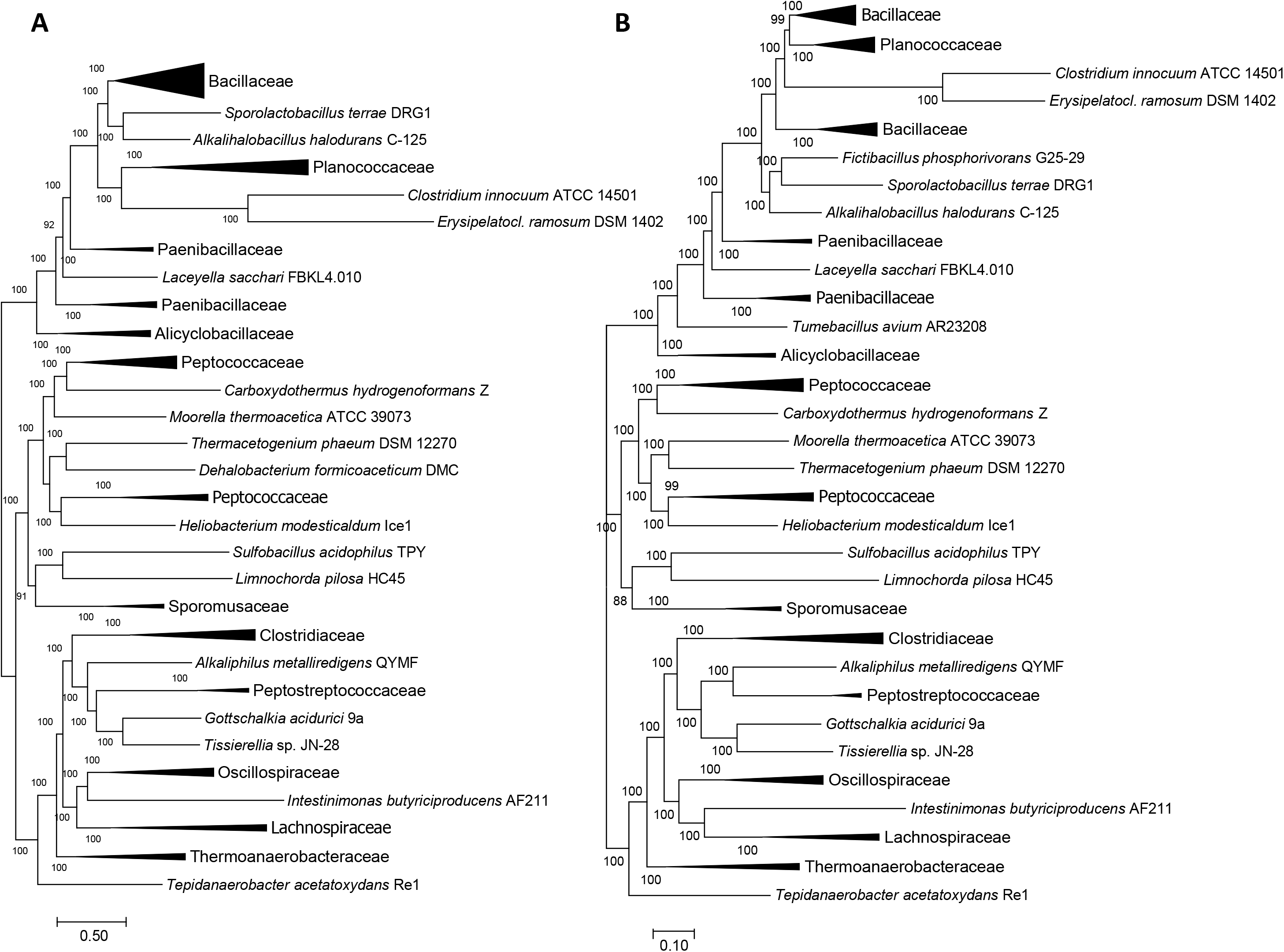
Phylogenic trees for sporulation and ribosomal proteins from Firmicutes. Maximum-likelihood phylogenetic trees were built using IQ-TREE2 (118) with the LG+F+R9 substitution model from concatenated alignments of 41 core sporulation proteins (A) and 54 ribosomal proteins (B) encoded in the genomes of 76 spore-forming members of the Firmicutes.

The evolutionary history of gains and losses of these sporulation genes (the same 41 except for *spoIIQ*) was reconstructed using GLOOME, by mapping the patterns of gene presence and absence in each lineage onto the phylogeny of 180 representative Firmicutes, to infer the posterior probabilities of gene presence in the ancestral tree nodes (121). This reconstruction placed all 40 core sporulation genes at the root of the Firmicute phylogenetic tree and suggested only a few gene gains (*jag* in three *Lactobacillus* spp, the dipicolinate transporter-encoding *spoVV*/*yjiH*/*ylbJ* genes in several non-spore-formers, *ycaP* in *Veillonella* and *Megasphaera*). An interesting example of a likely gain was the *spo0A* gene in *Macrococcus caseolyticus*, one of the few instances of this gene in non-spore-forming bacilli and the only such gene in the *Staphylococcaceae* members in our set (in GenBank, *spo0A* is also found in the members of the newly recognized genus *Mammaliicoccus*). The maximum-likelihood approach implemented in GLOOME considered gain of *spo0A* by *M. caseolyticus* more likely than the loss of this gene in five other staphylococcal genomes although the highly degraded sequence of this gene and its unusual placement in the phylogenetic tree of Spo0A (Figure S3) suggested the latter alternative as more realistic.

With all 40 core genes mapped to the root of the Firmicutes, GLOOME predicted extensive loss of sporulation genes in non-spore-forming lineages, such as lactobacilli, staphylococci, and listeria. A major loss of sporulation genes was also inferred for the asporogenous members of the order *Bacillales*, such as *Salimicrobium jeotgali*, *Gemella haemolysans*, and *Exiguobacterium* sp. (see Table S1). A similar picture was observed in the classes *Negativicutes* and *Tissierellia*. Many sporulation genes are conserved in the spore-formers *Methylomusa* and *Pelosinus* in the former class, and *Gottschalkia* and *Tissierella* in the latter, whereas asporogenous representatives of these classes retained very few sporulation genes (Table S1). Accordingly, GLOOME interpreted the absence of these genes in the non-spore-formers as lineage-specific gene losses. The two lineages mentioned above, *Planococcaceae* and *Erysipelotrichaceae*, showed loss of some sporulation genes in their spore-forming representatives and Spo0A-encoding non-spore formers, and a far more extensive loss in the Spo0A^−^ bacteria and non-spore-formers.

Among *Clostridia*, massive, family-wide gene loss was only seen in *Eubacteriaceae*, as most other families included both spore-forming and asporogenic genera. Here, dramatic differences in sporulation gene content were often observed within the same family. Thus, in the family *Peptostreptococcaceae*, the analyzed genome set included five members, two spore-formers, *C.difficile* and *Paeniclostridium sordellii*, and three non-spore-formers, *Acetoanaerobium sticklandii*, *Filifactor alocis*, and *Peptoclostridium acidaminophilum*(49). Accordingly, the first two encompassed near-complete sets of core sporulation genes (with the exception of those coding for the germination receptor GerABC and several others, see above), whereas the three non-spore formers showed the loss of at least 29 of the 40 analyzed genes (see Table S2). Further, many clostridial non-spore-formers still carried *spo0A* and retained a fair number of sporulation genes (Figure 1, Table S1). As an example, all six representatives of the order *Halanaerobiales* in the current set were non-spore-formers, but they all carried *spo0A,* and a major loss of sporulation genes was observed only in *Halanaerobium hydrogeniformans* (Table S1). This case, as well as the examples of lineage-specific gene loss in *Butyrivibrio proteoclasticus* (family *Lachnospiraceae*) or *Fastidiosipila sanguinis* and *Mageeibacillus indolicus* (family *Oscillospiraceae*) (Tables S1 and S2), show that dramatic loss of sporulation genes can occur on relatively short evolutionary distances.

## DISCUSSION

Sporulation is a complex process that involves dozens of genes that function in a tightly regulated fashion to ensure the survival of the bacterial cell under adverse conditions. Spores are resistant to a variety of damaging factors, including extreme conditions, and can persist for thousands – and potentially millions – of years (122). A recent study suggests that spores of *B. subtilis* could even survive on a simulated Martian surface (91). A better understanding of the sporulation mechanisms could help in finding new approaches to eradicate spore-forming human and animal pathogens. Furthermore, recent studies of the human microbiome have revealed a variety of spore-formers that remain to be characterized in detail (123), suggesting that the impact of Firmicute spore-formers on human health could be even greater than currently appreciated.

An essential step towards understanding the sporulation process is delineation of the set of genes that are necessary and, possibly, sufficient for sporulation across the entire diversity of bacterial spore-formers. Given that the functions of many genes involved in sporulation remain unknown, those genes that belong to the core set would become priority targets for structural and functional characterization. A major stumbling block for the identification of the core set of sporulation genes is the uncertainty of the sporulation status of many bacteria with sequenced genomes. While many genomes come from well-characterized type species, many others do not, and it is usually not known whether the sequenced strain exactly corresponds to the one that was described previously, in some cases, many years ago. A case in point is *Acetohalobium arabaticum*, which was initially described as a spore-former (124), but apparently lost the ability to sporulate during the laboratory passages (31, 125); we listed it as a non-spore-former (Table S1). Conversely, *Turicibacter sanguinis*, a Spo0A-encoding member of class *Erysipelotrichia*, was originally described as asporogenous (126) but subsequently shown to form typical subterminal spores (see Extended Data Fig. 4 in ref. (123)). The genome of *Turicibacter* sp. H121 included in this work has almost the same set of sporulation genes as *C. innocuum* and *E. ramosum* but lacks, among others, *divIVA*, *dpaB*, *spoIIGA*, *spoVK*, and *spoVN* (Table S2), making this strain unlikely to sporulate. The spore-forming representatives of *Erysipelotrichia* analyzed in this work, *C. innocuum* and *E. ramosum*, were occasionally described as forming very low numbers of spores (127, 128). Nevertheless, there was no doubt that these organisms are spore-formers, which made defining their set of sporulation genes a worthwhile exercise.

Previous studies by several independent groups (2, 32, 34–37) identified essentially the same set of ∼60 sporulation genes of *B. subtilis* that were shared by spore-forming members of *Bacilli* and *Clostridia*. However, in the genomes of some spore-formers, certain genes from this conserved core were found to be missing (32, 36, 37). The goal of this work was to reassess the distribution of the key sporulation genes using the widest possible selection of complete genomes from all branches of the Firmicutes. To this end, we employed the latest version of the COG database (39) and supplemented it with five recently sequenced genomes from various members of the *Erysipelotrichia*. This resulted in a set of 180 species from 160 genera, representing 45 different families from six classes of the Firmicutes. At this scale, most genera were represented only by a single genome of a single species, which provided a different perspective from most previous studies (2, 32, 34–37). Therefore, this work complements, rather than supersedes, those studies.

As discussed elsewhere, the COG database displays the patterns of presence-absence of orthologous genes (COG members) across all covered genomes and therefore allows straightforward identification of those genes that are missing in a given genome (39, 42). However, there is an important caveat: a single COG often includes several paralogous genes, which can give the impression that each of these paralogs has a wider phylogenetic distribution than it actually does. For example, *cwlJ* and *sleB*, members of COG3773 “Cell wall hydrolase CwlJ”, are both shown as universal (Table 1), although some genomes, such as *C. perfringens* or *C. difficile*, may encode only a single member of that family. Likewise, *dacB* and *dacF* are paralogous genes that encode closely related D-alanyl-D-alanine carboxypeptidases (penicillin-binding proteins 5* and 1, respectively, members of COG1686) that are involved in peptidoglycan metabolism. However, the former is expressed in the mother cell under SigE control, whereas the latter is expressed in the prespore under the control of SigF and/or SigG. Members of this COG, which also contains sporulation-independent paralog, *dacA*, are found in most firmicutes, both spore-forming and not.

Further, certain gaps in phyletic patterns can arise from errors in genome sequencing and annotation. In some cases, genes that were frameshifted in the examined genomes (Table S3) were found in full-size versions in the genomes of other strains or closely related species. Although we made an effort to address the potential annotation problems, fixing possible sequencing errors was not part of this project. As a result, we have recorded the instances of rarely missing genes in Table S6, but were mostly interested in cases where a certain gene was missing in more than one or two genomes, particularly when these genomes came from the same bacterial lineage.

In our previous work, we noted the absence of many key sporulation genes in the relatively large (4.8 Mb) genome of *Lysinibacillus sphaericus* C3-41 (129), but blamed it on the poor quality of the genome sequence, calling it “the worst offender” among all bacilli and excluding it from most analyses (32). Now, many of those genes were also found to be missing in the genome of the type strain of this species, *L. sphaericus* DSM 28^T^. Further, such genes as *spoIIB*, *spoIIM*, *spoIIIAA*, *spoIIIAB*, *spoIIIAD*, *spoIIIAF*, *spoVAA*, *spoVAB*, *spoVID*, *bofC*, *gerM*, *spmA*, *spmB*, and *yqfC* were missing in the genomes of all members of the family *Planococcaceae* (Table S4), to which *L. sphaericus* was recently reassigned (58). These observations showed that, rather than being “the worst offender” in terms of genome content, *L. sphaericus* could be a valuable model organism to study the basics of the engulfment process.

The expansion of the initial genome set to include two spore-forming members of *Erysipelotrichia*, *C. innocuum* and *E. ramosum*, provided a window into an even greater loss of core sporulation genes (Table S4). In these organisms, sporulation appears to be less frequent than in members of other lineages, making them somewhat similar to oligo-sporulating mutants of *B. subtilis*. Nevertheless, the ability of *C. innocuum* and *E. ramosum* to form spores in the absence of numerous core genes (Table S4) makes them useful model organisms for study minimal requirements for sporulation among the Firmicutes.

The conservation of core sporulation genes in *Bacilli* and *Clostridia* suggested that the ability to form spores was a common ancient feature of the Firmicutes (2, 4). Although the scenarios of the origin of Gram-negative (diderm) bacteria from spore-forming Gram-positive (monoderm) ones (130, 131) do not seem to be supported by phylogenetic trees of universal genes (120, 132, 133), understanding the fundamental mechanisms of sporulation and clarifying the origin of spore-formers remains a major goal in microbiology. Here, we used an expanded collection of the firmicute genomes to trace the phylogeny of the core sporulation genes within this phylum. Despite certain variability in the branching order (Supplemental file 2), proteins coming from the members of the same phylum typically formed well-supported clades. This was particularly clearly demonstrated by a comparison of the trees built from concatenated alignments of sporulation proteins and ribosomal proteins (Figure 3). We and others have previously demonstrated that such a ribosomal proteins-based tree is closely similar to the 16S rRNA gene-based tree and faithfully represents organismal phylogeny of the Firmicutes (28, 49, 134). The remarkable congruence of these trees strongly suggested vertical inheritance of the sporulation genes from a common ancestor of all Firmicutes. Further, in the Spo0A tree (Figure S3), proteins from non-spore-formers often grouped together with Spo0As from spore-formers of the same bacillar or clostridial family, suggesting that Spo0A-encoding non-spore-formers lost the ability to form spores, rather than acquired their *spo0A* genes. Accordingly, non-spore-forming Spo0A^+^ bacteria typically retain many more sporulation genes than Spo0A^−^ non-spore-formers (Figure 1). These observations lend further support to the hypothesis that the ability to form (endo)spores was an ancestral feature of the Firmicutes that emerged in this phylum at the early stages of bacterial evolution (4, 32). This ability remains a unique feature of some Firmicutes, distinguishing them from all other bacterial lineages. Evolution of Firmicutes from their spore-forming last common ancestor involved extensive, but in most cases, incomplete loss of sporulation genes in those lineages that lost the sporulation capacity. Subsets of the sporulation genes were also lost in several lineages of spore-formers. Experimental study of sporulation in these organisms with reduced complements of sporulation genes can be expected to yield further insight into the minimal biochemical requirements for sporulation.

Despite the substantial progress made in the past several years, a large fraction of sporulation genes remain uncharacterized. For some genes, this is reflected in their *y*-names, but many genes with specific names have been characterized only with respect to the timing of their expression, as being controlled by sporulation-specific sigma factors, or as encoding proteins of the spore coat, whereas their biochemical activities remain obscure. For example, spore maturation proteins SpmA and SpmB, which are conserved in all spore-formers except for the members of *Planococcaceae* (Table S6), participate in spore core dehydration and are required for heat resistance of spores (135, 136). However, the biochemical mechanisms of spore dehydration by SpmA and SpmB remain unknown. We hope that phylogenetic profiles of the key sporulation proteins derived in this work (Table S2) help identifying priority targets for experimental research and allow the selection of suitable model organisms for such studies.

## MATERIALS AND METHODS

### Genome coverage

The list of *Firmicute* genomes used in this work was taken from the recent release of the COG database (39). The COG genome set includes complete genomes of 175 members of the *Firmicutes*: 73 members of the class *Bacilli*, representing 59 different genera; 79 members of the class *Clostridia*, representing 76 genera; 10 members of the class *Negativicutes*, 9 members of the class *Tissierellia*, 2 members of the class *Erysipelotrichia*, 1 member of the class *Limnochordia*, and an additional genome of “*Ndongobacter massiliensis*”, which remained unclassified at the time of this work (137). Most genera were represented by a single genome, except for five members each of *Bacillus* spp. and *Streptococcus* spp., four members each of *Clostridium* spp. and *Lactobacillus* spp., three *Staphylococcus* spp., and two *Listeria* spp. The list of these organisms is available in Table S1 in the Supplemental material and also on the NCBI website https://ftp.ncbi.nih.gov/pub/COG/COG2020/data/cog-20.org.csv. Examination of the available publications allowed extracting sporulation characteristics for 173 out of these 175 organisms. For 74 of these, the ability to form spores was documented in the descriptions of the respective species (Table S1). Ninety-nine organisms were non-spore-forming, based on the explicit descriptions of the respective strains and/or their lineages (such as the order *Lactobacillales* and families *Acidaminococcaceae*, *Halanaerobiaceae*, *Listeriaceae*, *Peptoniphilaceae*, *Selenomonadaceae*, *Staphylococcaceae*, and *Veillonellaceae*). For two organisms, the sporulation status remained unknown, as their (in)ability to form spores had not been properly documented and the respective lineages included both spore-formers and asporogens (Table S1).

Since both members of the *Erysipelotrichia* included in the COGs were asporogenic, the genomic list for this class was supplemented with five recently sequenced complete genomes: two from spore-forming members of the genus *Erysipelatoclostridium*, [*Clostridium*] *innocuum* ATCC 14501 (Genbank accession <u>CP048838.1</u>) and *Erysipelatoclostridium ramosum* DSM 1402 (GenBank: <u>CP036346.1</u>), and three genomes of non-spore formers, *Amedibacterium intestinale* (GenBank: <u>AP019711.1</u>), *Faecalibaculum rodentium* (GenBank: <u>CP011391.1</u>), and *Intestinibaculum porci* (GenBank: <u>AP019309.1</u>). This brought the total set to 180 genomes, 76 of which came from proven spore-formers (Table S1).

### Protein selection

The list of sporulation proteins analyzed in this work was based on the lists of genes that were previously demonstrated to play a role in sporulation of *B. subtilis* and/or were regulated either by Spo0A or by sporulation-specific sigma subunits SigE, SigF, SigG, or SigK (2, 7, 12, 13, 15–18, 33, 34, 115, 138), as documented in the SubtiWiki database (38).

The selected genes were matched against the COG database (39) and the patterns of presence or absence of the respective COGs in the selected organisms were recorded in Table S2 in the Supplemental materials (in some cases, a single COG included several paralogous sporulation genes). The resulting set consisted of 237 protein families (COGs), which included 112 COGs for widely conserved genes from the previous COG releases, such as *ald* (*spoVN*), *ftsH* (*spoVK*), *ftsI* (*spoVE*), *ftsK* (*spoIIIE*), *spmA*, *spmB*, *spoVG*, *spoVS*, and so on (Table S2). That list was expanded by the addition of 125 sporulation COGs that were made available in the recent release of the COG database (39). While this list mostly contained genes (proteins) characterized in *B. subtilis*, it also included clostridial forms of SpoIIQ (COG5833) and small acid-soluble spore protein (SASP, COG5864), as well as the Amidase_6 domain (COG5877). The widely conserved sporulation genes (proteins) were assigned to one of the following eight functional groups: 1) Sporulation onset and checkpoints (22 genes); 2) Spo0A-regulated genes (9 genes); 3) Engulfment (12 genes); 4) Spore maturation, SigF- or SigG-regulated forespore expression (19 genes); 5) Spore maturation, SigE- or SigK-regulated mother cell expression (25 genes); 6) Spore cortex synthesis (9 genes; 7) Spore coat and crust proteins (8 genes), and 8) Germination proteins (8 genes) (see Table S2). Two more groups included sporulation genes conserved in Bacilli (51 genes) and narrowly represented genes (74 genes) (Table S2). This selection was based on the previous studies (2, 15, 16, 28, 32, 36, 37) and the data from the SubtiWiki database (http://subtiwiki.uni-goettingen.de/) (38).

### Verification of the COG profiles

The patterns of presence-absence of the sporulation genes in the 180 analyzed genomes were extracted from the COG master file cog-20.cog.csv, which is available in the https://ftp.ncbi.nih.gov/pub/COG/COG2020/data/ folder, matched against the assembly numbers listed in Table S1, and the counts of each gene in the 76 spore-former genomes were calculated, ignoring the paralogs. The cases of apparently essential sporulation proteins missing in certain spore-former genomes were verified using a recent version of the TBLASTn program (55) that allows selection of specific target organisms (or lineages) based on their entries in the NCBI Taxonomy database (56). Proteins from either *B. subtilis* reference set or closely related organisms (same genus or family) were used as queries. The resulting BLAST hits (cut-off E-value, 0.1) were verified using the CD-search (139) and compared against the protein sets in GenBank and RefSeq databases. Confirmed sporulation genes were used to correct the COG profiles. Newly translated sporulation ORFs were reported to GenBank and/or RefSeq.

### Phylogenetic analysis

Unless indicated otherwise, protein sequences for phylogenetic analyses were aligned with MUSCLE v5 (140), and the maximum-likelihood trees were inferred using a locally installed version of IQ-TREE 2 (118) with the best-fit model automatically selected by ModelFinder (119) and branch support calculated using Bayesian-like approximate likelihood test (141). For each COG that included paralogs, a consensus was produced as described in (145); a pairwise similarity score between each sequence and the consensus sequence was calculated using the BLOSUM62 matrix (with the value of -4 assigned to a gap in the sequence, matching a non-gap character of the consensus sequence); the paralog with the maximum score was used in the alignment. Individual phylogenetic trees, constructed using IQ-TREE 2 (118), are presented in Supplementary file 2.

For the phylogenetic analysis of Spo0A, the alignment included sequences from all 118 Spo0A-encoding organisms in the analyzed set: 76 spore-formers, 40 non-spore-formers, and 2 organisms with uncertain sporulation status (Table S1). The MUSCLE alignment was edited to remove variable-length linkers between the REC and DNA-binding domains (corresponding to the residues 131-148 in the Spo0A of *B. subtilis*), which left 225 informative sites. The maximum-likelihood tree of Spo0A (Figure S3) was constructed with IQ-TREE 2 (118) using the LG+R6 substitution model.

The SpoIVCA tree (Fig. S4B) was constructed from an alignment of 1,220 SpoIVCA-like site-specific DNA recombinases/integrases from bacteria and phages that were identified in a PSI-BLAST (55) search. The alignment contained 476 informative positions. The approximately maximum-likelihood tree was built with FastTree 2 (142) using “–gamma” option to rescale the branch lengths and the WAG (143) model of amino acid evolution.

For phylogenetic analysis of sporulation, alignments of 41 core sporulation proteins from 76 spore-formers were constructed [the 40 genes marked in Table 1 with the addition of YmfB (COG0740), a SASP-degrading paralog of ClpP (144)]. The few missing proteins (Table S2) were replaced with the respective sequences from closely related organisms (the same genus, where available). For the overall phylogenetic tree of sporulation core, these 41 alignments were concatenated, resulting in an alignment with a total of 10,129 informative sites, and the maximum-likelihood tree (Figure 3A) was built using IQ-TREE 2 (118) with the LG+F+R9 substitution model. The ribosomal proteins-based tree (Figure 3B) was built from a concatenated sequence alignment of 54 ribosomal proteins from 76 spore-formers with IQ-TREE 2 and the LG+R8 substitution model.

## ACKNOWLEDGEMENTS

This study was supported by the Intramural Research Program of the U.S. National Library of Medicine at the National Institutes of Health.

