## Supplemental Figures S1 to S8 and Tables S3 to S85 for "Conservation and evolution of the sporulation gene set in diverse members of the Firmicutes"

**Table S1 (Excel). Properties of 180 organisms with complete genomes used in this work**
**Table S2 (Excel). Presence of 237 sporulation genes (COGs) in selected genomes**

**Supplemental file 1. Figures S1-S8, Tables S1-S8.**

Figure S1. Genome and proteome sizes of spore-formers and non-spore-formers
Figure S2. Patterns of the presence of 237 sporulation genes (COGs) in selected genomes
Figure S3. Phylogenetic tree of Spo0A proteins
Figure S4. SpoIVCA-like site-specific recombinases/integrases from bacterial *skin* elements.
Figure S5. Structure of the SpoIIQ–SpoIIIA complex from *B. subtilis* and the loss of its components in the six members of *Planococcaceae* (A) and *Erysipelotrichaceae* (B).
Figure S6. Variability of *spoIIQ* genomic neighborhoods within the *Bacillaceae*
Figure S7. Sequence alignment of small acid-soluble spore proteins from *Erysipelotrichia*
Figure S8. Maximum-likelihood phylogenetic tree of 180 firmicutes built from a concatenated alignment of 54 ribosomal proteins

Table S3. Frameshifts, nonsense mutations, and insertions in core sporulation genes
Table S4. Widespread sporulation genes missing in certain organisms (by organism)
Table S5. Conservation of the previously defined sporulation core
Table S6. Core sporulation genes missing in certain genomes (by gene)
Table S7. Split *sigK* genes in diverse firmicutes
Table S8. COG-based annotation of uncharacterized sporulation protein

**Supplemental file 2. Phylogenetic trees in Newick format (text file)**

76 spore-formers sporulation protein tree (Figure 3A)
76 spore-formers ribosomal protein tree (Figure 3B)
Spo0A_tree (Figure S3)
Ribosomal protein tree for 180 organisms (Figure S8)
Trees for individual sporulation proteins (41 trees)

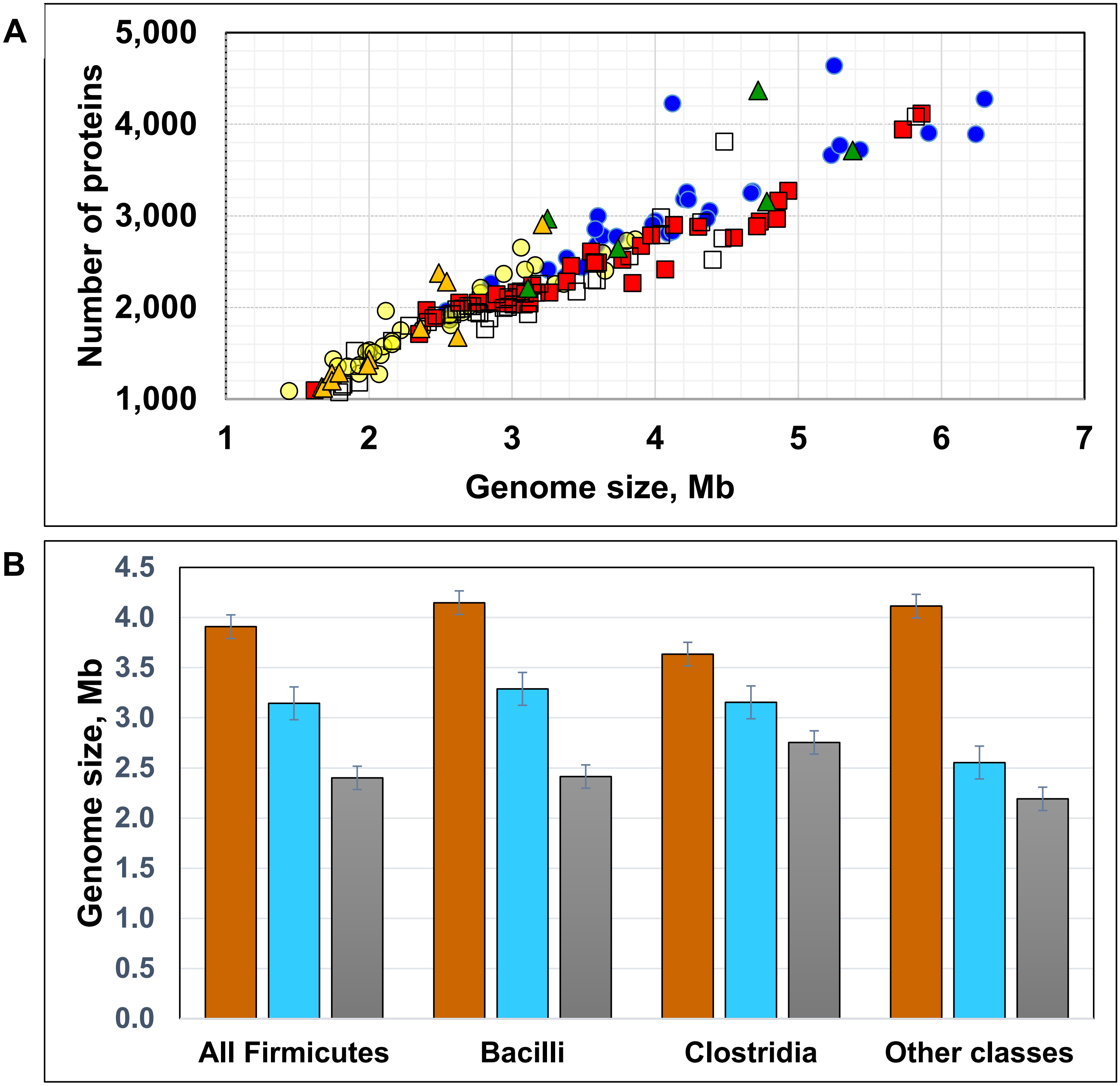


**Figure S1. Firmicute spore-formers generally have larger genome sizes than asporogens.**

**A. Genome and proteome sizes of the 180 organisms in the analyzed set.** The symbols indicate the following organisms: blue circles, spore formers of the class *Bacilli*; yellow circles, bacillar asporogens; red squares clostridial spore-formers; empty squares, clostridial asporogens; green triangles, spore-formers from other classes; orange triangles, asporogens from other classes.

**B. Comparison of the genome sizes** in spore-formers (left columns, brown), Spo0A-encoding non-spore-formers (central columns, cyan), and *spo0A*⁻ non-spore formers (right columns, gray).

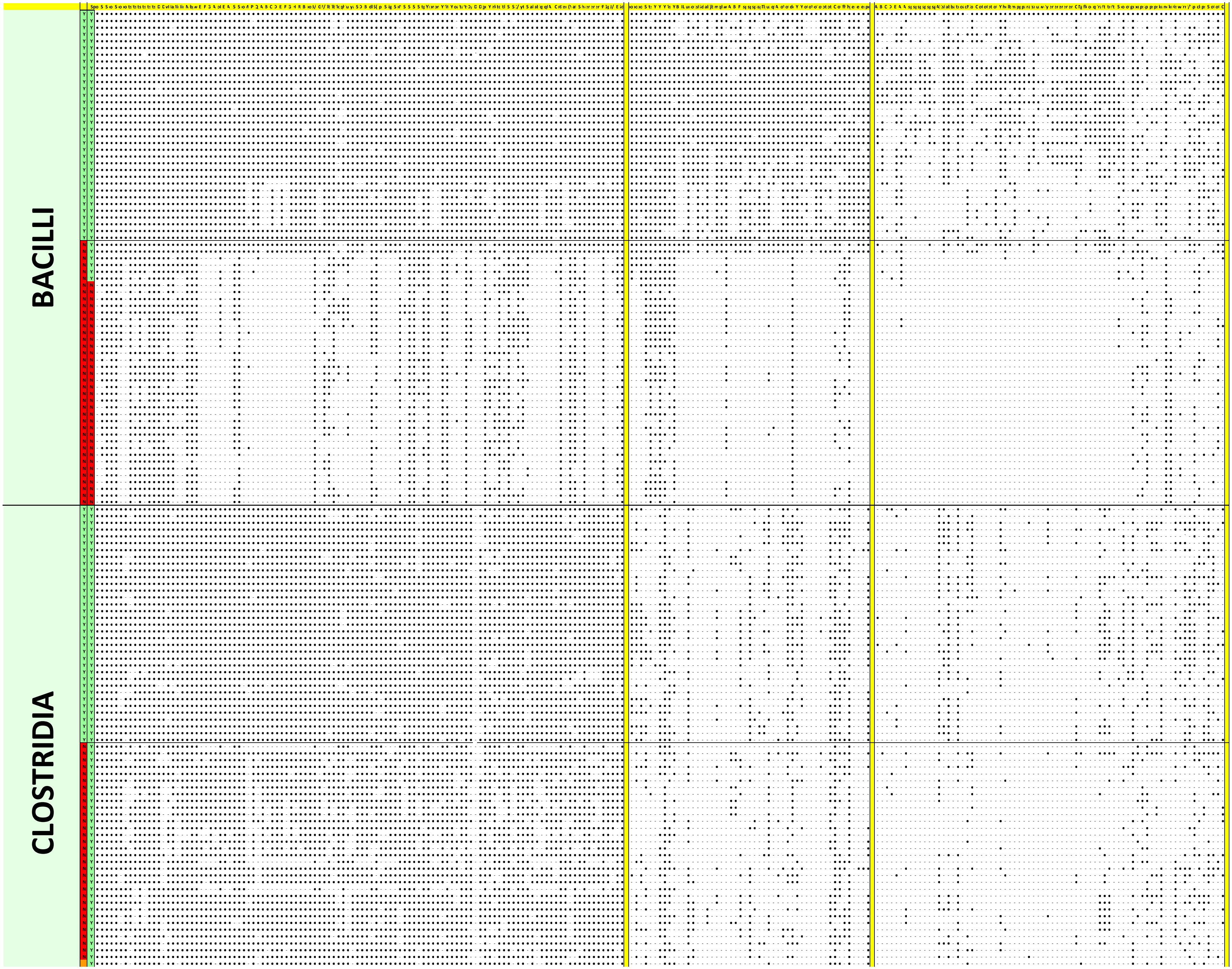
BACILLI
CLOSTRIDIA

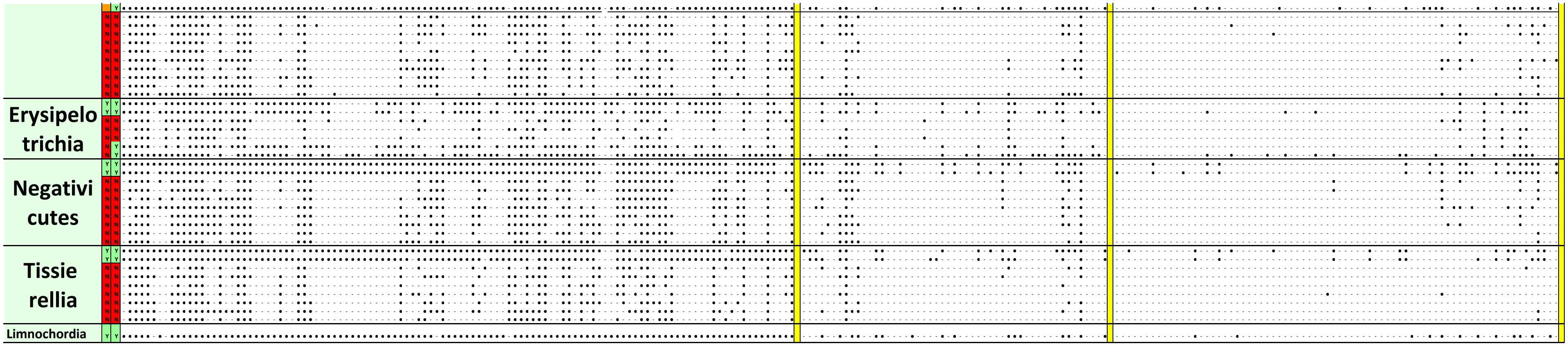


**Figure S2. Patterns of presence of 237 sporulation genes (COGs) in selected genomes**.
The vertical blocks represent a) widely conserved sporulation proteins; b) those conserved mostly in *Bacilli*, and c) narrowly conserved sporulation proteins. The horizontal blocks represent six classes of Firmicutes, sorted by their ability (Y on the green background in the first narrow column) or inability (N on the red background in that column) to form spores. The second narrow column shows the presence (Y on the green background) or absence (N on the red background) of the *spo0A* gene in the organism's genome. See Table S2 for details.

### Figure S3. Maximal-likelihood phylogenetic tree of Spo0As from 118 members of Firmicutes

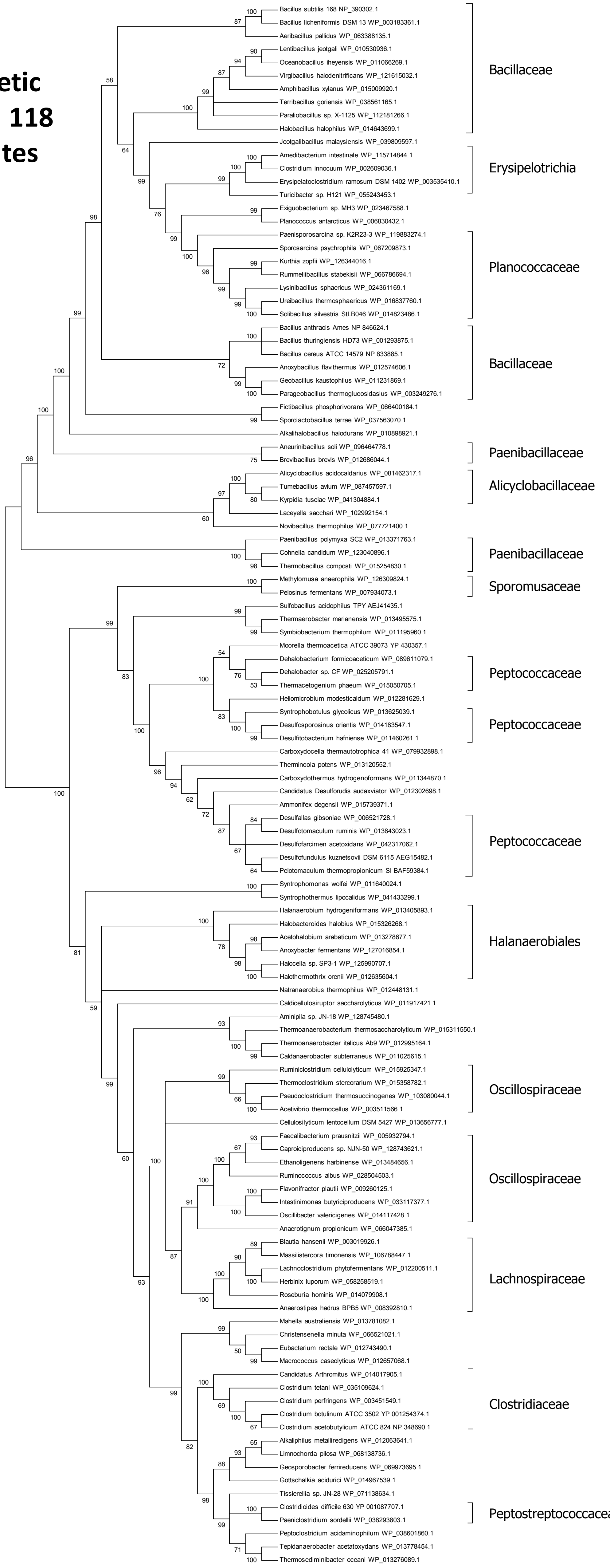

**A**

```
Sec_Struct          --EEEEEEEE----------HHHHHHHHHHH-----EEEEEE----------HHHHHHHHHHHH----EEEEEE--------HHHHHHHHHHH----EEEEEE-
Bs_SpoIVCA      1   -MIAIYVRVSTEEQAIKGSSIDSQIEACIKKAGTKDVLKYADEGFSGELLERPALNRLREDASKGLISQVICYDPDRLSRKLMNQLIIDDELRKRNIPLIFVNG  103
Aflv_0803       1   -MKAIYARVSTEESAQKGYSIAHQIEEAKKKAKTDDVMIYTDEGFSGEFLERPGLERLRNDIRNGMIDTVIVYDPDRLARNLMHQLLLDDEFRKYNVELLFVNG  103
BBR47_19300     1   -MIGIYARVSTEEQAKSGFSLDDQLRECRKKAATSEAAEYVDDV-SGEFLDRPALSRLRRDAKEGLITKIVCLDPDRLSRKLMNQLLITDEFDKRGIELVFVNG  102
B0W44_05430     1   -MIGIYARVSTEEQARTGFSLQDQIKECKKKARTNEVIEYIDDGFSGEFLDRPALTKLRNDVKEGIISKVICLDPDRLSRKLMNQLIITDEFDRLDVELIFVNG  103
Amet_2475       1   MNIAIYSRKSKFTG--QGESVENQITICREYIEK5EINVYEDEGFSGGDTDRPQFQRLIKEARAKRFDVLICYRLDRISRNISNFARTIEMLENNNISFVSVKE  106
EQM14_04150     7   TRIAIYSRKSRFTG--KGESIENQIELCRQYIAQ7SLLVYEDEGFSGGSTDRPQFKKMMADARGKKFSILVCYRLDRISRNIGDFARLIEELGSLSISFVSIRE  113
CD630_12310     1   MSVAIYLRKSRADEE-AEKQGEFETLSRHKSTLL9VIEIKEELVSGESIIHRPKMLELLKEVEENKYDAVLVMDLDRLGRG2KDQGIILETFKESKTKIITPRK  112
CTC_01067       2   NKICIYLRKSRADEELEKTLGEGETLSKHRKALL9IVEIKEEIVSGESLFFRPKMLELLKEVENKQYTGVLVMDMQRLGRG2QDQGIILETFKKSNTKIITPMK  114
Nther_1738      1   -MIGIYARVSTEEQAKTGTSIQDQLRACKEYLSK6QLQEYIDEGISGEIFERPALSKLRNDSERGNLTAVICFDPDRLSRKLMHQLIITEELNEHGVELLFVNG  108
PTH_1125        1   MRAAVYVRVSTEDQARHGYSLQEQKEACRCRAVD4TVLEFADEGVSGATLDRPGLQGLRELIRSGQIDLVVVRDPDRLSRKLSHQLILTEEIEKAGVRLEFLDF  107
Ccel_0430      28   LEVGGYIRISTKKDSQITSIENQKKYIMEWASVN3IVDFYIDMKTGAYSYLRQEMVRMKNDISSGKIKGIVSKEISRTSRD2LDIIELKRSLANQGAFFISIKE  133
TPY_3324        1   MKAAIYLRVSSDEQALHGYSLAAQRHACRERARQ4EIQEWADEGIPGTVWDRPGLTQLRQAVSARAIDLVVIYDPDRFARSLALQLVVTEEIERAGVQLEFVNF  107
Tph_c16790      1   MRAAIYARVSTEEQAVHGFSLAEQVSACKKKAVE4EILEFADEGISGATLNRPGLEALREAVREGKIDCLILRDPDRLSRRLAHQLLLTEEFEKAGVKLEFLDF  107
JBW_01921      10   EPSCFYLRKSREDQE-AEARGDGETLAKHKKALF9ITKIFEEVVSGESIIHRPEMLNLLKEIELGKWKSVFCMDIDRLGRG2QDQGLIIETFKQAQTKIVTPRK  121
Curi_c13490     1   MKAVIYSRKSKFTGKGESIENQVQMCKEYAYNHF5DITVYEDEGFSGGTTDRPQFQLMLKDAKQKKFDVLICYRLDRISRNIADFAVLIEELQNYNVSFVSIRE  108
4BQQ_A          3   TYAGAYDRQSRERENSSAASPATQRSANEDKAADLQREVERDG16AFGTAERPEFERILNECRAGRLNMIIVYDVSRFSRL2MDAIPIVSELLALGVTIVSTQE  121

Sec_Struct          -----------HHHHHHHHHHHHHHHHHHHHHHHHHHHHHHH--------------------------HHHHHHHHHHHHH-------HHHHHHH----------
Bs_SpoIVCA    104   EYANSPEGQLFFAMRGAISEFEKAKIKERTSSGRLQKMKKGMIIKDSKLYGYKFVKEKRTLEILEEEAKIIRMIFNYFTDHKSPFFGRVNGIALHLTQMGVKTK  207
Aflv_0803     104   EYANTPEGKLFFSMRGAISEFEKAKIRERTMAGRKRKAKEGKVVKNDKIYGYDYDKKTGQLIVNESEAEIVRFIFDSFTQGR---FKGINGIALHLTERNIPTK  204
BBR47_1930    103   EYAKTPEGQLFYSMRGAIAEFEKAKINERMSRGRREKARQGRVLRDFQIYGYSYDSKKEQIVINEAEAAVVRLVFDLFTQPN-ELVQGMNGIAVYLTNKGVPTK  205
B0W44_05430   104   EYAKTAEGQLFYSMRGAIAEFEKAKINERMSRGRREKARQGRVVRDYQIYGYDYDKETEQFIINESEAEIVRLIFDLFTKPN-DLVRGINGIAKYLTEKGVPTK  206
Amet_2475     108   FDTATPMGRAMLYIASVFAQLERETIAERIRDNMLQLARTGRWLGGNTPTGFES13KMFKLSPIDEELTLVTLLFDKYLEFK-----SLSKLETYLIQNDLTTK  217
EQM14_04150   115   FDTASPMGRAMMYIASVFSQLERETTAERIRDNMRELAKTGRWLGGTTPTGYRS13RAFRLVFQPDEIKTVRLIFQKFLETD-----SLTQVETFLIQNSIPSK  225
CD630_12310   113   TYDLTDEFDEEYSEFEAFMARKELKLISRRMQRGRIKSV2GNFIGTSAPFGYDAV2GRKERILVPNKDADVVRTIFDLYINE---DMGCSKISKYLNNLGIKTA  215
CTC_01067     115   TYDLSDDFDEEYTEFEAFMSRKELKMINRRMQGGRVRSV2GNYIATNPPLGYDIH2KKSRTLKINPHECEIIKLIFKLYTEG----NGAGSIAEHLNNLGYKTK  216
Nther_1738    109   YYQKTPEGNLFYSLRGAISEFEKAKITERMKRGRLEKARQGSVLRDYNVYGYNYNKNNRSLEINKKESEIVKLIFDLFTRPE-GQNLGINGIANYLTKKGIPTK  211
PTH_1125      108   DWKDTPDGRLFYAIRGAIAEFEKEKIRERMARGKTQKAKQGGMPIGFYNYGYVYEPETGKVRLHETEAKVVEEIFKWFVQED----IGINGVAKRLNEAEVPSR  207
Ccel_0430     134   GYDSRTDDDEFLLVIHAGLAQKERKVTGSRVKITQMIKA2GKTNVPTPALGYKLSPDGQHIIINPATAEIYQLIVKKFLEGW-----GRLKICKYLNSQGIRTN  233
TPY_3324      108   DWQDTPEGKLFYAMRGAFSEYEREKIRLRTSLGRLQKARQGHMPMAVAPYGYRYDPATARLEVHPQEAPIVVRMFTECVQEQ----KSLNSIARQLTTEKIPTR  207
Tph_c16790    108   EWKDTPEGRLFYSIKGAIAEYEREKIRERMTRGKLQKARQGGIPVNFDVYGYHYDPATGRVSINEEEAATVSCIFNWFTTED----IGIAGVANRLNEMGIPTK  203
JBW_01921     122   IYDLNNEWDEEYTEFESFMARKELKIITRRLQGGRIRSL2GNYLGAVPPYGYLIE3RKRYLIKHPEQCEPTSLIWKAYQLDM-----GANKIANELNQRGYLSY  223
Curi_c13490   110   QFDTSTPMGRAMMYIASVFAQLERETIAERIRDNLQQLA2GRWLGGITPTGFIS13KAYKLSTVQEEVELVKLLYNKYIEIK-----SLTGLETYCLRNNIKSK  219
4KIS_A          1   --------------------------RDRMVMGKIKRIEAGLPL3KGRTFGYDVIDTKLYINEEEAKQLRLIYDIFEEEQSI-----TFLQKRLKKLGFKVRTY   75

Sec_Struct          ---------------------------------EEEEEE---------------------------HHHHHHHHHHHH--------------------------
Bs_SpoIVCA    208   KGAKVWHRQVVRQILMNSSYKGEHRQYKYDTEGSYVSKQAGNKSIIKIRPEEEQITVTIPAIVPAEQWDYAQELLGQSKRKHLSISPHNYLLSGLVRCGKCGNT  311
Aflv_0803     205   MGKSVWHRQVVRQILMNETYVGKKPQNKWNTEGMLANKYRNEKVPMRLRNKDEWIYVDVPAIVSAEQFQQAQMLLEQARRRYTKASKRQYLLSGLIRCAQCGNT  308
BBR47_1930    206   RGASVWHRQVVRQMLMNEAYVGRFYQNKWNTEGMLGNQ3PDEKVRMKMRPKDEWISLPCPSIIDEVKFEHAQRLLKESRRRWAGRSFNEYLLSGLVRCGSCGNT  311
B0W44_05430   207   RGANVWHRQVVRQMLMNRAYIGKFYQNRWNTEGMLGNK3KDERIPMKERPREEWIPVPCPAIIDRQTFDHAQNLLSESRRRWTKQSKRRYLLSGLLRCAKCGNT  312
Amet_2475     218   -TGKCFQKHSLRTILTNPVYAQTDGALYAYFSSNGAQ21VKKGKSVKRKDMSEWIVAI3KGIIDGREWIQVQHMMHINKDK2RQGTSHTALLSGLIRCAGCGSF  342
EQM14_04150   226   -TGKRYTRFAIKNILENPVYMAADPDAYRYFAGSGAE21QQHGKANRAREMSNWIVAI3QGILSGEDWIRVQKMLERNKSK2RKPKNNTALLSGLLRCGRCGSY  350
CD630_12310   216   -TGANWYNSAITNIIKNKVYCGYIQWQK--KDYKKSKN-PNKIKTVKLRPKDEWIEAK3EPLISEITWKKAQNILKKNGHV-SYGNQIKNPLAGIVICKNC---  313
CTC_01067     217   -FNNNFSRSSVLFILKNPIYIGKVTWKK--KEIKKSKN-PNKVKDTRTRDKSEWIVVD3EPMISMKMWNKAQEILNNKYHI2QLVNGPANPLAGIVICSKC---  316
Nther_1738    212   TNKQTWHRQVVKQILQNPVYIGSFYQNKWKAEATRDSP3ATKSISPKVRDQSDWIEIKCPAIITESKFYKAQELLKQSRRKWVKQGKRLYLLSGLVKCGLCNNT  317
PTH_1125      208   -KGKRWHKQVVRQVLVNPVYKGTWQY---------------------------KDICIPVPAIIDEAVWLKAQEKIRGARRLWAGQRKHDYLLSGIVTCGECGQT  284
Ccel_0430     234   RGNASWSTNSIYAILTNPVYLGITMYNITLTVRDDTG-----KAKRMVRPRDQWIVKE3QPLITKEKFERIQQLIQDKKQK3EWSCTKKYLLSGLVYCGSCGSK  336
TPY_3324      208   KGLGVWHRQVIRQILQNPVYTGTFYANRYETYGVGLNR3AGTKVRRKVRPRSEWIGIPAPRLIDPVIWEAAQQRIRERTR----TPGTSYLLAGLVRCGLCRQT  311
Tph_c16790    208   RGAGCWHRQVVKQILVNPVYKGEWKYGKVDWH------------TRTPRPAEEVITIPVPAIVDYRTWSGAQEKLQRIRRLWSKKGRHQYLLSGLLVCADCSNT  299
JBW_01921     224   -SGKKWSASSVLAILKNPAYAGVNAWKK--VASKKSTTRTGRETRHRPREEQIWIYDCHEPYITLEEFEQVQDMLSKKYHP2QLLNGVTNPLAGLIKCDIC---  322
Curi_c13490   220   -NGANFYVHTLKIILENPVYATADKKLYNYLIGNGHA21QVKGTTDRNRDQSDWIISV3EGIISSDMWIKANEIMEMNRDK3KRYINTNALLSGILICENCGSF  345
4KIS_A         76   NRYNNW--------LTNDLYCGYVSYKD----------------------KVHVKGIHEPIISEEQFYRVQEIFSRMGKNPNMNKESASLLNNLVVCSKCGLG  148

Sec_Struct          -EEEEE------EEEEEE------------------HHHHHHHHHHHHHHHHHHHHHHHHHHHHHHHHHHHHHHHHHHHHHHHHHHHHHHHHHHHHHH--
Bs_SpoIVCA    312   MTGKKRKSHGKDYYVYTCRKNYSGAKDRGCGKEMSENKLNRHVWGEIFKFITNPQKYVSFKEAEQSNHLSDELELIEKEIEKTKKGRKRLLTLISLSDDDD  412
Aflv_0803     309   MTGRWMKNWGKYHREYTDRKNTAGAKHRGCGMVVRADEIEQEVWDYLLDLFNHPERIQEFQ3DAENSYEKKELERLEKEIDRARKGRKRLLSLVTMSDDDE  411
BBR47_1930    312   MSGRKVKNWSQHIFEYTDVKNTAGAKNKGCGRRVRCEQLDNQVWETVRSWLNNPDEIAIAV3KVETPFEQVELERLQKELEKTKAGRKRLLKLFASVEED-  412
B0W44_05430   313   MTGRKHRNWGADVFIYTDEKNYSGAKHPGCGMRVRCEKLDKDVWDKISHWLNNPDEIATVE3TDEVDFEQAEIIRLETEIKKAKEGRKRLLNLFAEGLE--  412
Amet_2475     346   KYGQVSKETGKRIYYYVCNNKNI-SRGVKCKCPNLQGMKVDKGIMEELKVLLANDVLKSLD3NVNINTKKLEVKELQNKIHNKHKAIDNLVKQLSKSDDEF  446
EQM14_04150   354   KLTRRKNRDGEPVYTYLCELKEK-SRGRKCSVPNINGNSLDQAVCSEIISLPEDGSAFLQH8KEDRDDLDTELERLNQSIQNAEKAISNLVSALAEGRSS-  459
CD630_12310   314   -GRPLVYRPYADHDYIICYHPGCNKSSRFEFIEAAILKSLEDTVKKYQLKASDIDLDRNNK3IEFQKRVLKGLETELKELGKQKNKLYDLLERGIYDE---  412
CTC_01067     317   -KFKMVMRKLKGVDRLLCRNNKCDNISNR---YDSTEKAIIQALERYLKEYRINISNQNKT2IKPYERQVNILEKELAALNEQKLKLFDFLERGIYDE---  411
Nther_1738    318   MTGRRVKNWGEYVFIYTDKKNNP-GIYQGCG-LNIPCSELDKVVSEVVFNWLTDLNEIKSQ5EGQNKHIELDISETESKLSKTKQQKKKLLDLIGEDT---  417
PTH_1125      285   MTGVYSKWWNKKDRRYTCFKGYQGARHRGCL3YVLAGYVESAVWEQVKDWLQDPGALASEA3SPRVEDYLKELERTEKLLSEVEKGREAVLDALASGLFE-  387
Ccel_0430     345   KTAKLKIESERTPKDYYYYYFDR-KTSGVCR--NSKANFHMDIVEQKVIEKVKEILLSYDQ8YLFDTRYIKEKKENE-RLRDRLQTVNSAIRREQSAYEAD  449
TPY_3324      312   LVGRRDQSWGRPNRIYTCRKHSAGAAHPGCR2ERLDADALEDAIWQGMKSYGSRLLQ------QLAERPAGEGNDRHARLDRITRAQGHLLDALAEGLG--  403
Tph_c16790    300   MGGSFIKWWGTGRRRYTCRRAGCVSRSFGCR2KAIPADVLENAVWEQIKGFLCDAGAIARE4MSKKSTTEREYENIKGRLKAIEKGKEKIVDALASGLL--  402
JBW_01921     323   -GGSMIYRPYQHQQFPHLMCYNRNCINKSCR-FEYVEKKLLEGLQLWLNDYRTQWDIFIPK5VNSKEKVHQNLKKELKELEQQKDNLHDLLEKGIYNAATY  425
Curi_c13490   348   KGATRVNKDKSIAYYYLCELKEK-SRGQNCNIKNIRGDKLDKLVIEELKRLSSQESRLHSK8ETSQSNIESEIDILKKNIESNNKSIANLINALAQGEG--  452
4KIS_A        153   RKDTVSRGKKYHYRYYSCKTYKHTHELEKCG----NKIWRADKLEELIIDRVNNYSFASRN--IDKEDELDSLNEKLKIEHAKKKRLFDLYINGSYEVSEL  247

Sec_Struct          ---HHHHHHHHHHHHHHHHHHHHHHHHHHHHHHHHH----HHHH-HHHHHHHHHHHH------HHHHHHHHHHHHHHEEEE----EEEEE-
Bs_SpoIVCA    413   LDIDEIKAQIIELQKKQNQLTEKCNEIQSKMKVLDDTSSSENALKRAIDYFQSIGADNLTLEDKKTIVNFIVKEVTIVDSDTIYIETY  500
Aflv_0803     412   LDLEEVKEEIRNLQQREKELIEQYNQLEQQLKEIQESKPNEHILQAAIDFFLNHK-EQLDFEIKQTLIRKVVKEIRVIDKENIEIHLF  498
BBR47_1930    413   ISEEDIRQGLKELKEKEDKLNQRSNELHEQAKLQIANEYNRNLIQEAADYYLSKAQDELSFGDKKELIRHVVREVRVYEESVEIYTF-  499
B0W44_05430   413   IGQEEIRESLKGLKRKEEHLTEELEELKKRYSELSNKQYSANLIKEAAAYYLNHGKEELSFDDKQALIRYVVKEIRVYKDGIEIYTF-  499
Amet_2475     447   -VSDYILNEIKRLIQANKVLSQELENDHKEIQLL----NIDLLKMPSVGFSEIVDGTLYESKRNFINGIIEKIVWDGESLDIHLLNI-  528
EQM14_04150   460   AASRYIIRQIDEWKEKEDSLEAGIQQLKDRAEIRF-LPDEQIEMAADALRSFSSAFGSMTVDEKRCALRSLI-DRVVWDGETVSICLF  545
CD630_12310   413   DTFIERSNNISSRTEEIKDSIKTVKNRLSTVKKDN-SKIIEDIKTVLSLYHDSDSLGKNKLLKSVINKAVYYKSKEQKLDSFELMVHL  499
CTC_01067     412   NTFLERSKNIEERITKTSSGIEKINDIINKEKKVV-KEEDVIKFQKLLDGYKNTDDIKLKNELMKKLVNKVEYTKNKRGDTFGIDIFP  498
Nther_1738    418   ISIQEAREYLQSCNQKEKKLSQKLKELQAHKKNIKKQYQSKQSKHIISSTFAKIKNYNEEDLINQQQIIRTMIREIIIYKNSIVINTH  505
PTH_1125      388   LDARTKAKLADLKRRKERLEQRKKELVLTVRGASG--AVARMDELRALASEVLGRIDELDFSEKKALVRALVAQVVVSGKGKRGANGL  474
Ccel_0430     451   ITLDEYKTRLDELRRQKLSLDEKLEKINSTPAQSG--SLEVANKLKETIRSLHNLEQDFQYEIIRKLINKIYIKDNYSLEFEFTFSD-  535
TPY_3324      404   --------DPAVIRRRLRALEEERRALDPSLSPPS--VRSGRVFEWEAIPVYWRRE--ILEVLVRRIVVGDGQLTIVLKTDPENISVA  479
Tph_c16790    403   ELDERTKNKLADLKSRWERLENRKKELERHLLAAG--DGAAALEELQRLALRLLERIDDLSFEEKRALIRAVVSQIIVAGRPQPGKAE  488
JBW_01921     431   KLSERITTVQKNLDKAAFALSKEIQQEESRKNIIP-KIEKVLKTYEKSNDPTEKNKFLKSILEHATYRKEKWQKGDTFTLVLRPRLPR  517
Curi_c13490   453   SLAAKHILDQINKLESENANLKEKLLDLRESIDDKHSLEENFEIMNRMINIFKNTIDDAEDDEKRSLIRNLVDRITWDGENIDIVMFG  540
4KIS_A        252   NDIDAQINYYEAQIEANEELKKNKKIQENLADL 284
```

**B**

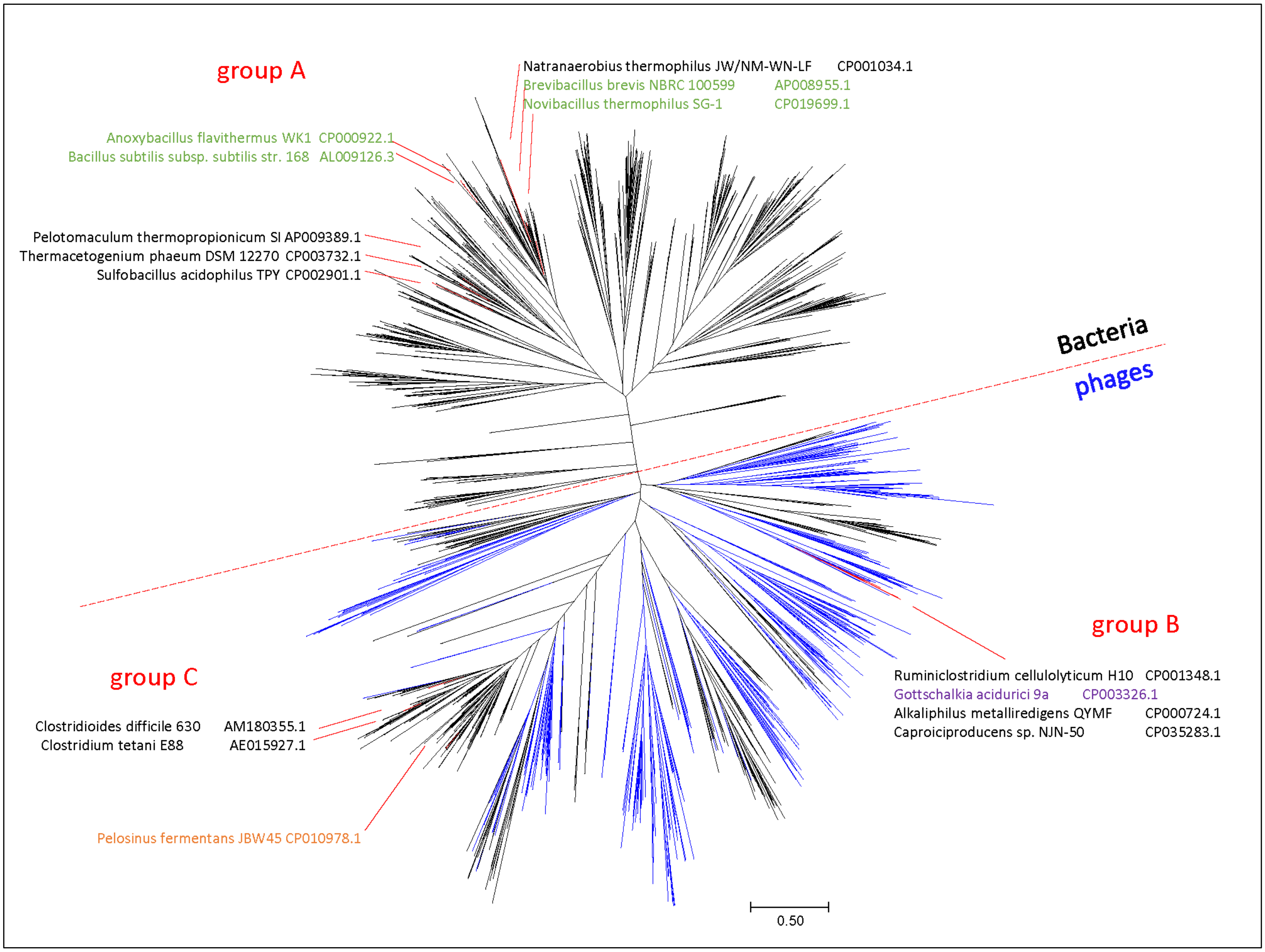


**Figure S4. SpoIVCA-like site-specific recombinases/integrases from bacterial *skin* elements.**
**A. Sequence alignment**. Proteins are listed under their genomic locus tags as shown in Table S7, their names in the first block are linked to the respective entries in the NCBI protein database. The top line shows secondary structure prediction by Jpred (http://www.compbio.dundee.ac.uk/jpred/, Drozdetskiy *et al*., 2015): H, α-helix, E, β-strand. The bottom line shows aligned sequences of serine recombinase from *Streptomyces* temperate phage phiC31 (GenBank: CAA07153, PDB: 4BQQ) and *Listeria innocua* prophage integrase (GenBank: CAC97653, PDB: 4KIS), see Rutherford *et al*. (2013) *Nucleic Acids Res.* 41: 8341–8356, PMID: 23821671). Conserved residues are shown in bold, conserved hydrophobic residues are shaded yellow.

**B. Maximum-likelihood phylogenetic tree of bacterial SpoIVCAs and their phage homologs**. The tree was built using IQtree from an alignment of 1,220 SpoIVCA proteins from bacteria and phages, identified in a BLAST search. Proteins from *skin* elements are indicated by the source genome name and genomic entry in GenBank (green indicates members of *Bacilli*, black – *Clostridia*, salmon – *Negativicutes*, and purple – *Tissierellia*. The respective branches are shown in red.

**C (below). Genomic maps of *skin* elements showing the locations of *spoIVCA* genes**. *Skin* elements are split into two groups, A and B+C, according to the phylogeny of their SpoIVCAs (panel B). *spoIVCA* genes (labeled SR) are shown in red, *spoIVCB* genes (labeled sigKn) and *spoIIIC* genes (labeled sigKc) are in brown with red borders. Genes of known phage-related proteins are colored, white shapes indicate uncharacterized ORFs

C

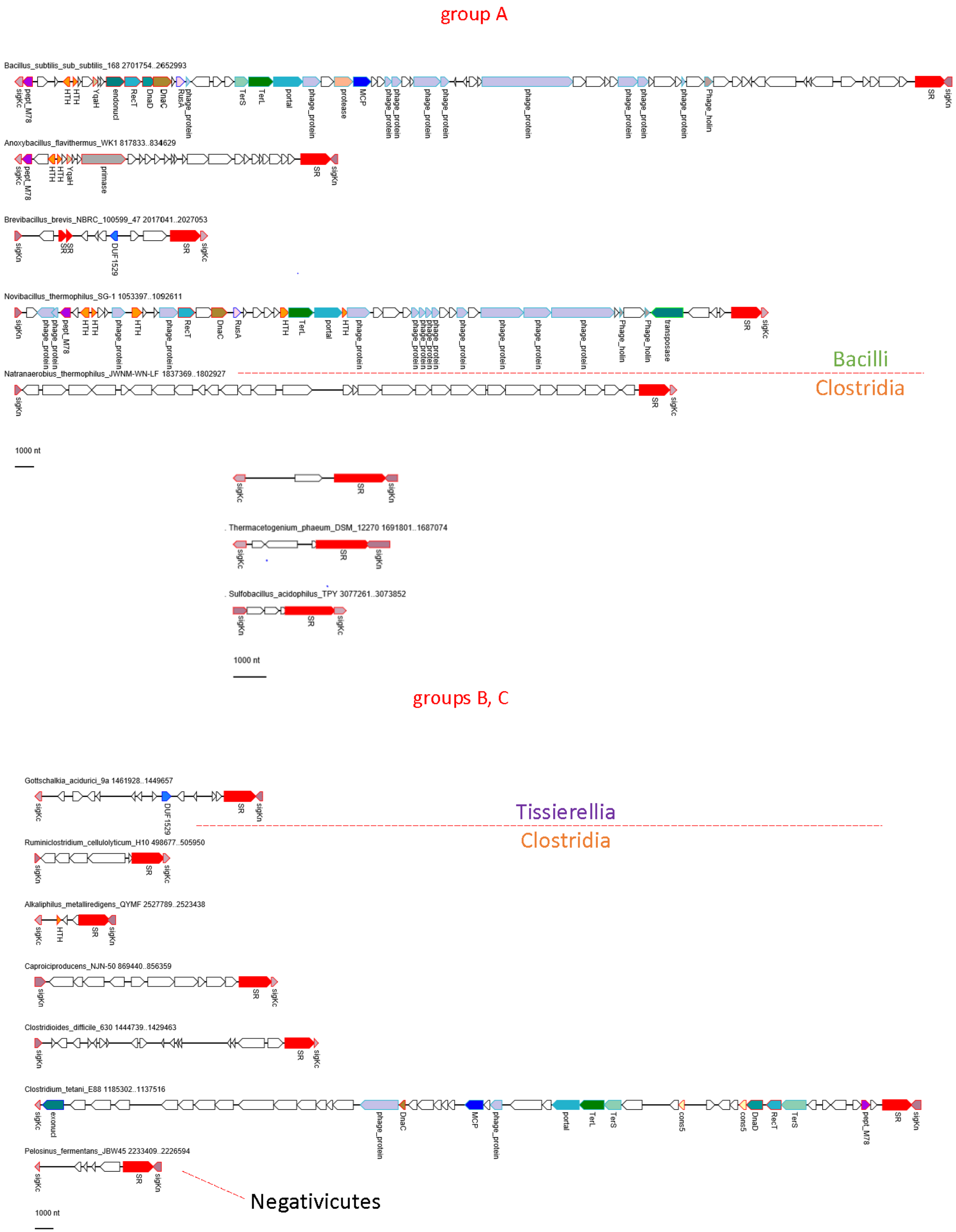
group A
Bacillus_subtilis_sub_subtilis_168 2701754..2652993
Anoxybacillus_flavithermus_WK1 817833..834629
Brevibacillus_brevis_NBRC_100599_47 2017041..2027053
Novibacillus_thermophilus_SG-1 1053397..1092611
Natranaerobius_thermophilus_JWNM-WN-LF 1837369..1802927
Bacilli
Clostridia
1000 nt
Thermacetogenium_phaeum_DSM_12270 1691801..1687074
Sulfobacillus_acidophilus_TPY 3077261..3073852
1000 nt
groups B, C
Gottschalkia_acidurici_9a 1461928..1449657
Tissierellia
Clostridia
Ruminiclostridium_cellulolyticum_H10 498677..505950
Alkaliphilus_metalliredigens_QYMF 2527789..2523438
Caproiciproducens_NJN-50 869440..856359
Clostridioides_difficile_630 1444739..1429463
Clostridium_tetani_E88 1185302..1137516
Pelosinus_fermentans_JBW45 2233409..2226594
Negativicutes
1000 nt

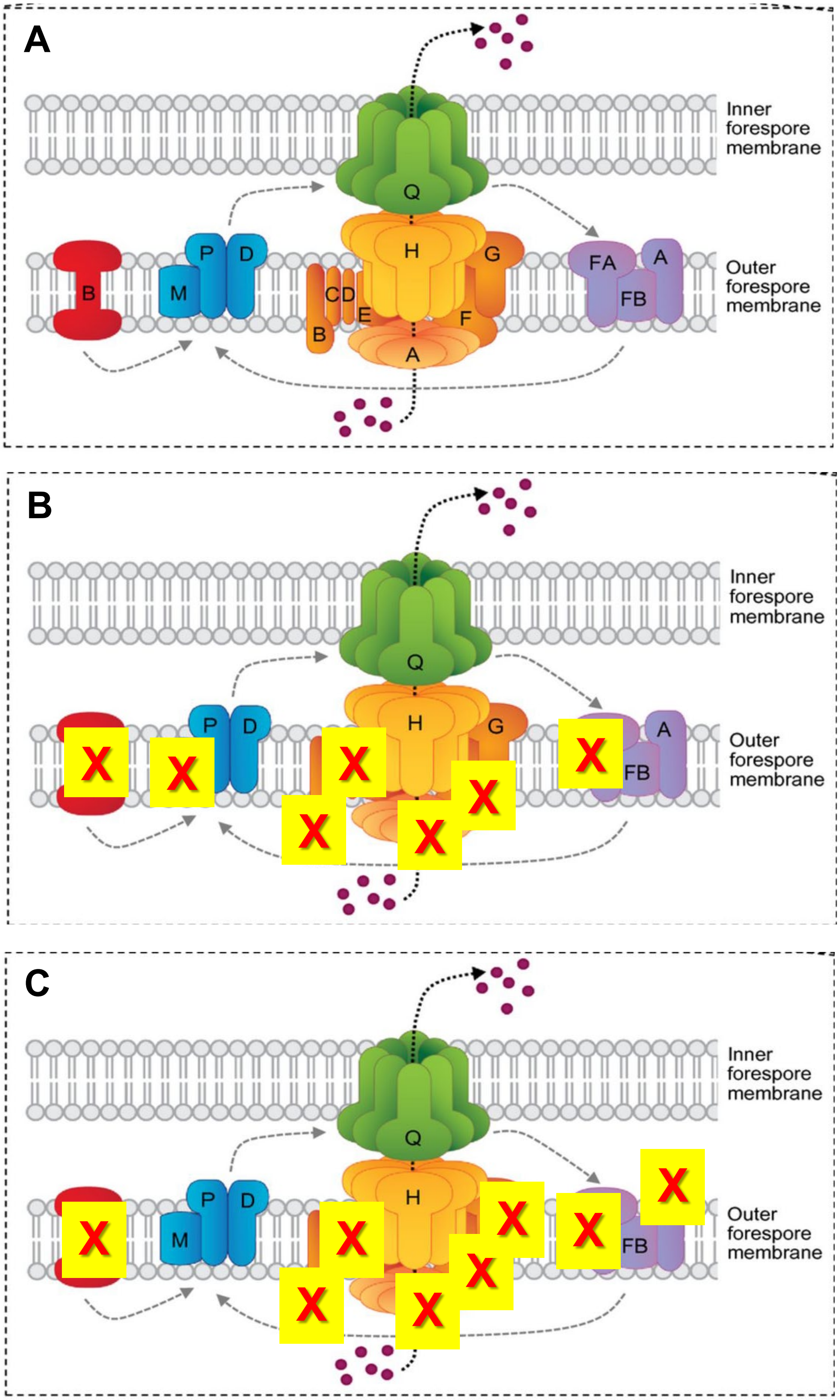


**Figure S5. Loss of the components of engulfment complex in Planococcaceae and *Erysipelotrichaceae*. A.** Organization of the engulfement complex (taken with permission from Crawshaw *et al*. (2014) *FEMS Microbiol Lett,* 358:129–136). **B.** In *Planococcaceae*, this complex shows the loss of *spoIIIAA, spoIIIAB, spoIIIAD, spoIIIAF, spoIIB, spoIIM*, and *spoIVFA.* **C.** Genomes of *Erysipelotrichaceae* members show additional loss of *spoIIAG* and *bofA*, while *spoIIM* is present.

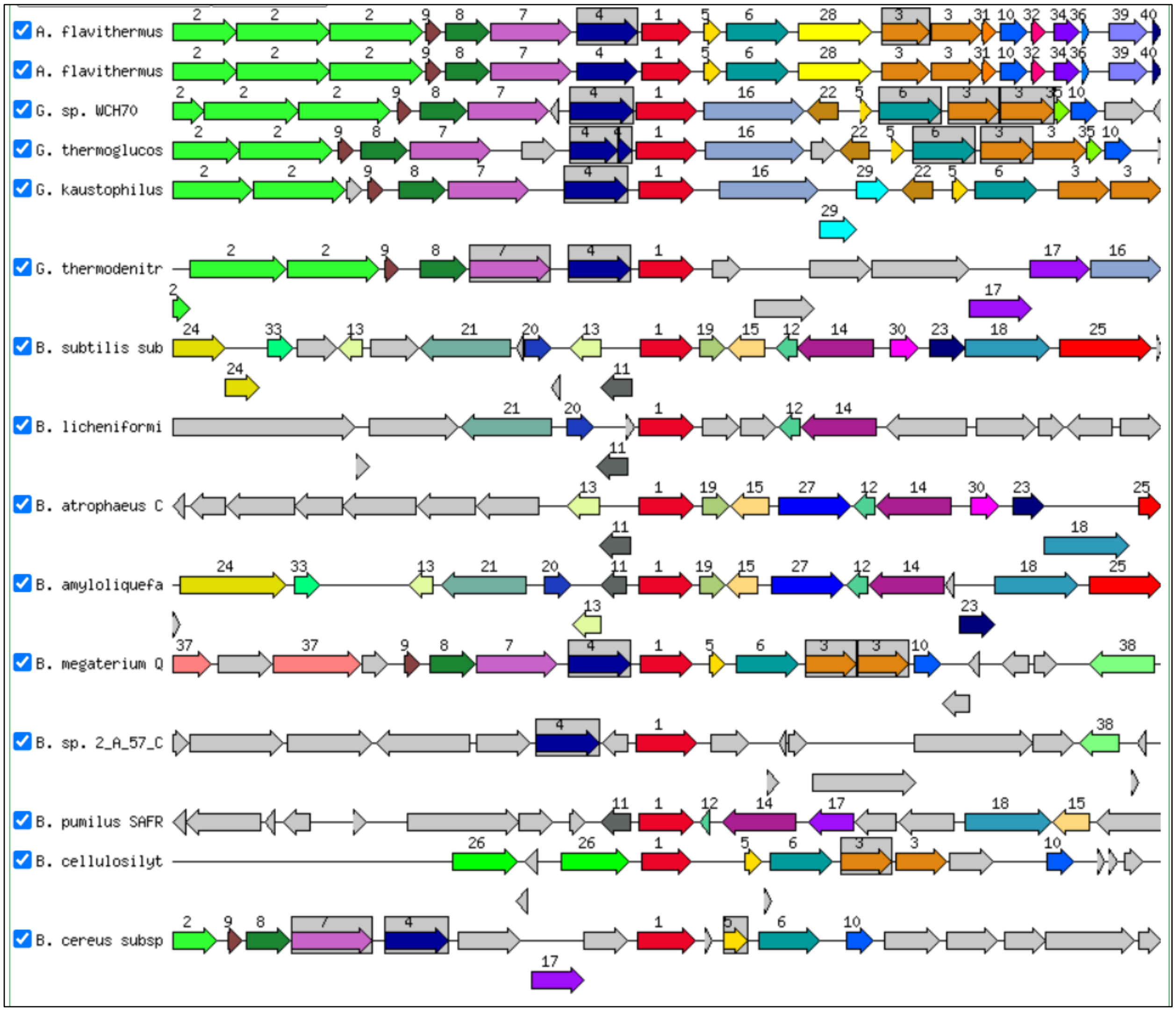


**Figure S6. Genomic neighborhoods of the *spoIIQ* gene in Bacillaceae.**

A SEED (http://pubseed.theseed.org/, Overbeek *et al*., Nucleic Acids Res. 2014, 42, D206-D214) rendering of the genomic neighborhoods of the *spoIIQ* gene (in red) in various members of Bacillaceae. The same neighborhood as in *B. subtilis* is seen in *B. amyloliquefaciens* and *B. atrophaeus* but not even is other members of the genus *Bacillus*. Most common neighbors of *spoIIQ* are, *spoIID* (no. 4, dark blue), spoIIID (no. 5, tan), *mbl* (no.6, sea green), *murA* (no. 7, purple), and *spoIIT* (*ywmB*, no. 8, dark green).

```
Sec_struct          ----------------HHHHHHHHHHHHHHH----------HHHHHHHHHHHHHHHHHHHHH-----------
SSPA_BACSU      2   ANNNSGNSNNLLVPGAAQAIDQMKLEIASEFGVNLGADTTSRANGSVGGEITKRLVSFAQQNMGGGQF-----   69
SSPB_BACSU      1   -MANQNSSNDLLVPGAAQAIDQMKLEIASEFGVNLGADTTSRANGSVGGEITKRLVSFAQQQMGGRVQ-----   67
SSPC_BACSU      5   SRSRSNNNNDLLIPQAASAIEQMKLEIASEFGVQLGAETTSRANGSVGGEITKRLVRLAQQNMGGQFH-----   72
SSPD_BACSU      1   ----MASRNKLVVPGVEQALDQFKLEVAQEFGVNLGSDTVARANGSVGGEMTKRLVQQAQSQLNGTTK-----   64
```

*Erysipelatoclostridium ramosum* DSM 1402

```
EDS17625.1      1   MSQSNSSRNKLVVPGAKNAIDQMKYEIANEFGVNLGPDTTARDNGSVGGEITKRLVAMGQAQMSSSNKYNQSK   73
EDS17626.1      1   MSQNNSSRNKLVVPGAQNAIDQMKYEIANEFGVNLGPDTTARANGSVGGEITKRLVEMGQSQMSSSNNYNQSK   73
```

[*Clostridium*] *cocleatum*

```
NDO41902.1      1   MSQNNSSRNKLVVPGAQNAIDQMKYEIANEFGVNLGPDATSRANGSVGGEITKRLVAMGQSQMSSS-NYNQSK   72
NDO41905.1      1   MSQNNSSRNKLVVPGAQNAIDQMKYEIANEFGVNLGPDTTARENGSVGGEITKRLVEMGQKQMTSSSRYNQSK   73
```

[*Clostridium*] *spiroforme* DSM 1552

```
EDS75125.1      5   -MSQYSSKNKLVVPGAQNAIDQMKYEIANEFGVNLGPDTTARENGSVGGEITKRLVAMGQSQMSSS-KYNQSK   71
EDS75126.1      1   -MSQNSNRNKLVVPGAKNAIDQMKYEIANELGVNLGPDASARSNGSVGGEITKRLVEMGQKQMSASSNYNQSK   72
```

*Turicibacter* sp. H121

```
AMC09182.1      3   NNSNNSSSNKLLVPGAQNAIDQMKYEIASEQGVQLGADATARQNGSVGGEITKRLVKQAQQQLGS----QNQQ   71
AMC09183.1      1   ----MSRRNKLVVPGAQNAIDQMKYEIAREFGVNLGPDTSSRQNGSVGGEITKRLVAMAQQQLGG----TSQK   65
AMC09184.1      1   ----MATRNKLIVPGAQEAVDQLKMEIANELGVNFGRKATAEENGAVGGEITKRLIAQAKQSL--------IK   61
AMC09185.1      1   -MANQSGRNKLVVPGAENAINQMKAEIANEFGVHLGPDATARQNGSVGGEITKRLVAMAQQQLG-------SK   65
AMC09186.1      1   ----MASRNKLVVPGAQNAIDQMKAEIAHEFGVTLGPDTSARQNGSVGGEITKRLVKQAQQQLGS----QNQQ   65
AMC08965.1      1   -MSNQSNSNKLVVPGAQAAIDNMKAEIANEFGVQLGADATSRQNGSVGGEITKRLVAMAQQQLG------QTK   66
AMC08966.1      1   -MSNQSNSNKLVVPGAQAAIDNMKAEIANEFGVQLGADTTARQNGSVGGEITKRLVAMAQQQLG------QTK   66
PF00269         1   ----------LLVPEAKEALDQFKYEVANELGVPLG6DLTSRQNGSVGGEMVKKMIEMAEQQL----------   58
```

**Figure S7. Sequence alignment of small acid-soluble spore proteins from *Erysipelotrichia*.**
The upper block shows sequences of four SASPs from *B. subtilis* and the secondary structure of its SspC (GenBank: AAB92494, PDB: 2Z3X). Protein names are linked to the respective entries in the NCBI protein database. Conserved residues are shown in bold, turn residues shaded green, hydrophobic residues shaded yellow.

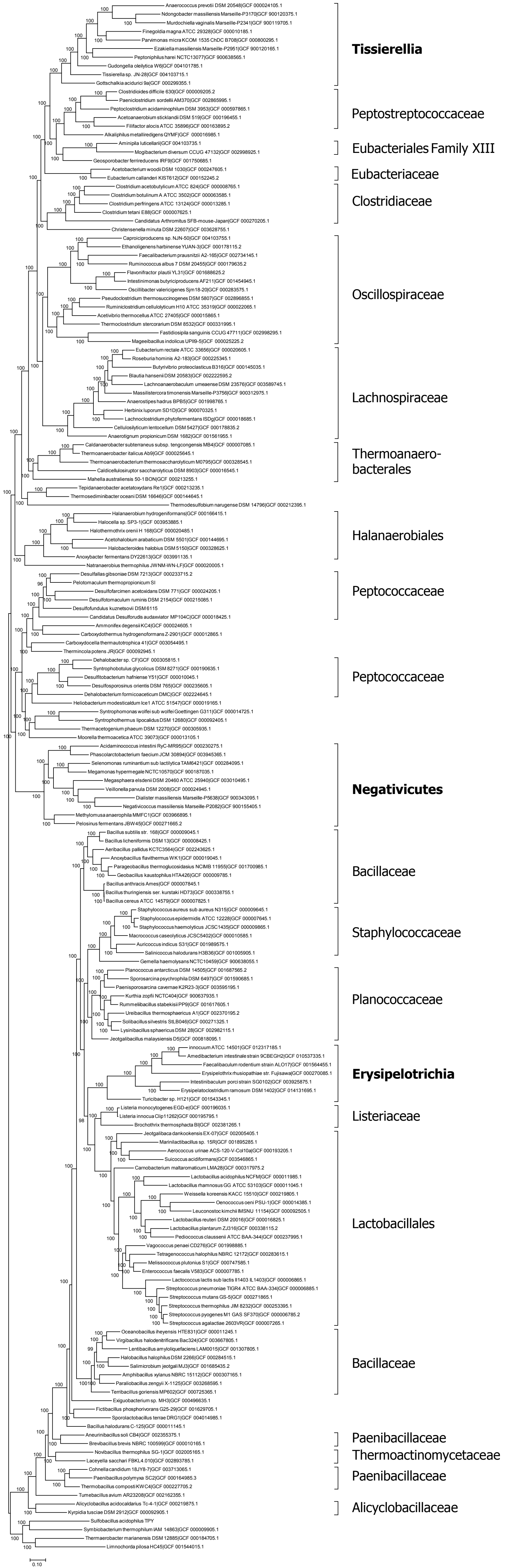


**Figure S8. Ribosomal proteins-based tree of all 180 members of Firmicutes**

**Table S3. Frameshifts, nonsense mutations, and *skin* element insertions in core sporulation genes**

| Organism name, GenBank accession no. of genome sequence | Protein name | COG no. | Genome locus tag(s) | First ORF frame, boundaries, protein ID (if available) | Second ORF frame, boundaries, protein ID (if available) |
|---|---|---|---|---|---|
| **Bacilli** | | | | | |
| *Alicyclobacillus acidocaldarius* subsp. *acidocaldarius* Tc-4-1, CP002902.1 | YlbJ | COG3314 | TC41_1554, TC41_1555 | Frame +2: 1528823..1529038, AEJ43485.1 | Frame +1: 1529038..1530165, AEJ43486.1 |
| *Bacillus cereus* ATCC 14579, AE016877.1 | SpoVK | COG0464 | BC_3710 (split into 3 ORFs) | Frame -3: 3675288 to 3675590, | Frame -1: 3675062 to 3675316. |
| | | | | Frame -2: 3674635 to 3675081 | |
| *Bacillus thuringiensis* serovar *kurstaki* str. HD73, CP004069.1 | SpoIIIAH | COG5828 | HD73_4489, HD73_4488 | Frame -1: 4330237..4329974, AGE80067.1 | Frame -2: 4329972..4329595, AGE80066.1 |
| *Jeotgalibacillus malaysiensis* D5, CP009416.1 | SpoIVA | COG5831 | JMA_19950, JMA_19940 | Frame -3: 1854201..1854437, AJD91312.1 | Frame -3: 1852959..1854197, AJD91311.1 |
| | GerAA | COG5091 | JMA_20360, JMA_20350 | Frame -3: 1889157..1889900 AJD91353.1 | Frame -3: 1888602..1889153, AJD91352.1, stop codon at 1889154 |
| | SpoIIM | COG1300 | JMA_20410 | Frame -1: 1895102..1895303, AJD91358.1 | Frame -3: 1894782..1895048 |
| *Lentibacillus amyloliquefaciens* LAM0015, CP013862 | DpaA | COG5842 | AOX59_02005 | Frame +1: 412924..413448 | Frame +3: 413448..413786 |
| | SpoVAEB | COG5839 | AOX59_01030 | Frame -2: 209795..210049 | Frame -1: 209703..209796 |
| *Novibacillus thermophilus* SG-1, CP019699.1 | DpaA | COG5842 | B0W44_14890 | Frame -3: 3056674..3056919 | Frame -2: 3056024..3056689 |
| *Oceanobacillus iheyensis* HTE831, BA000028.3 | CotJC | COG3546 | OB1667, OB1666 | Frame -3: 1709834..1710175, BAC13623.1 | Frame -3: 1709348..1709840, BAC13622.1, stop codon at 1709834 |
| *Paenibacillus polymyxa* SC2, CP002213.2 | SpmB | COG0700 | PPSC2_14790 | Frame -3: 3336263..3336698, ADO57119.1 | Frame -3: 3336704.. 3336793 |
| *Paraliobacillus zengyii* X-1125, NZ_CP029797.1 | SpoIVB | COG5832 | DM447_RS11010 (split into 3 ORFs) | Frame -2: 2170800.. 2171285 | Frame -3: 2170547..2170810 |
| | | | | Frame -1: 2170000..2170551 | |
| *Sporolactobacillus terrae* DRG1, CP025689.1 | YaaT | COG1774 | C0679_00245 | Frame +3: 52053..52661 | Frame +2: 52520..52879 |
| *Ureibacillus thermosphaericus* A1, NZ_AP018335.1 | SpoIIIAG | COG5828 | UtA1_RS10560 | Frame +2: 2194205..2194615, missing start codon | |

| | | | | | |
|---|---|---|---|---|---|
| **Clostridia** | | | | | |
| *Ammonifex degensii* KC4, CP001785.1 | SigK | COG5814 | Adeg_1441 | Frame +3: 1446546..1446620 | Frame +1: 1446628..1447170 |
| *Candidatus* Arthromitus sp. SFB-mouse-Japan DNA, AP012202.1 | YrbC | COG0217 | SFBM_0872 | Frame -3: 941010..940995 | Frame -1: 940290..940871 |
| *Heliobacterium modesticaldum* Ice1, CP000930.2 | SpoIIIAA | COG3854 | HM1_0273, HM1_0274 | Frame: -1: 508377..508958 ABZ82892.1 | Frame: -2: 507860..508369 ABZ82893.1 |
| **Negativicutes** | | | | | |
| *Methylomusa anaerophila*, AP018449.1 | YlbJ | COG3314 | MAMMFC1_03949 | Frame +1: 4255171..4256010, BBB93240.1 | Frame +2: 4256009..4256239 |
| **Interrupted genes** | | | | | |
| *Laceyella sacchari* FBKL4.010, CP025943.1 | CotJC | COG3546 | C1X05_12780 | Frame +3: 2604060..2604290 | Frame +1: 2612416..2612766 |
| | | | C1X05_12825 | Interrupted by a 8.1-kb *skin*-like element; SpoIVCA= C1X05_12810 , AUS09617.1 (517 aa), coded by 2609180..2610733 (+). | |
| *Ruminiclostridium cellulolyticum* H10, CP001348.1 | SpoIIID | COG5830 | Ccel_3110, Ccel_3179 | Frame +3: 3645999..3646124 | Frame +3: 3710868..3711011 |
| | | | | Interrupted by a 64.7-kb *skin*-like element; SpoIVCA=Ccel_3111, ACL77402.1 (511 aa), coded by 3646232..3647767 (-). | |

#### Table S4. Core sporulation genes missing in selected genomes (by organism)

| Organism name, GenBank accession no. of the genome | Genome size, Mb | Missing widespread sporulation genes |
|---|---|---|
| ***Bacilli*** | | |
| *Alicyclobacillus acidocaldarius subsp. acidocaldarius* Tc-4-1, CP001727.1 | 3.12 | *spoIIB, spoIIP, spoIVFA, spoVV, degV, germM, sapB, ytlD* |
| *Jeotgalibacillus malaysiensis* D5, CP009416.1 | 4.12 | *spoIIIAB, spoIIIAC, spoIIIAD, spoIIIAE, spoIIIAF, spoIVFA, spoVAC, spoVAD, cotJC, gerAB, gerM, gpr, ykuD, yheC, ypeB, yyaD/ykvI* |
| *Lysinibacillus sphaericus* DSM 28, CP019980.1 | 4.68 | *spoIIB, spoIIM, spoIIIAA, spoIIIAB, spoIIIAD, spoIIIAF, spoIVFA, bofC, gerM, spmA, spmB, gerW/ytfJ, ykuD/yciB, yqfC* |
| *Paenisporosarcina* sp. K2R23-3, CP032418.1 | 2.54 | *spoIIB, spoIIM, spoIIIAA, spoIIIAB, spoIIIAD, spoIIIAF, spoIVFA, spoVK, bofA, bofC, germ, gerW/ytfJ, sleL/yaaH/ydhD*, *spmA, spmB, ykuD/yciB, yqfC*, *ytaF*, yyaD/ykvI, |
| *Rummeliibacillus stabekisii* PP9, CP014806.1 + CP014807.1 | 3.42 | *spoIIB, spoIIM, spoIIIAA, spoIIIAB, spoIIIAD, spoIIIAF, spoIVFA, spoVK, bofC, gerM, gerW/ytfJ, spmA, spmB, ykuD/yciB, yqfC* |
| *Solibacillus silvestris* StLB046, AP012157.1 + AP012158.1 | 3.98 | *spoIIB, spoIIM, spoIIIAA, spoIIIAB, spoIIIAD, spoIIIAF, spoIVFA, spoVK, bofC, gerM, gerW* (*ytfJ*), *spmA, spmB, yqfC* |
| *Sporosarcina psychrophila* DSM 6497, CP014616.1 | 4.67 | *spoIIB, spoIIIAA, spoIIIAB, spoIIIAD, spoIIIAF, spoIVFA, bofC, gerM, gerW/ytfJ, spmA, spmB, yqfC* |
| *Ureibacillus thermosphaericus* A1, AP018335.1 | 3.49 | *spoIIB, spoIIM, spoIIIAA, spoIIIAB, spoIIIAD, spoIIIAF, spoIVFA, spoVK, bofC, gerM, spmA, spmB, ykuD/yciB, yqfC* |
| ***Clostridia*** | | |
| *Candidatus* Arthromitus sp. SFB-mouse-Japan, AP012202.1 | 1.62 | *bofA*, *cwlJ/sleB*, *gerM*, *jag*, *sapB*, *sleL/ydhD*, *ykuD*/*yciB*, *ypeB*, *ytlD*, *yunB*, *yyaC* |
| *Sulfobacillus acidophilus* TPY, CP003179.1 | 3.55 | *spoIID, spoIIIAB, bofA, cotJC, degV/yviA, divIB, gerM, jag, sapB, yaaT/ricT, ylbF/ricF, ypeB, yunB,* |
| *Cellulosilyticum lentocellum* DSM 5427, CP002582.1 | 4.71 | *spoIIM, spoVS, spoIVFB, cwlJ/sleB, ylmC/ymxH, ypeB, yunB* |
| ***Erysipelotrichia*** | | |
| [*Clostridium*] *innocuum* ATCC 14501, CP048838.1 | 4.72 | *spoIIR*, *spoIIIAA, spoIIIAB, spoIIIAC, spoIIIAD, spoIIIAE, spoIIIAF, spoIIIAG, spoIIID, spoIVFA, spoVS, spoVID, spoVIF, bofA, bofC, cotQ, cwlJ/sleB, gerAB, gerAC, gerD*, *gerW/ytfJ, lipC, minC, ricF(ylbF), safA, sleL/yaaH/ydhD, sspA, yabQ, ykuD/yciB, ymfJ, ypeB, yrkC yyaC, yyaD* |
| *Erysipelatoclostridium ramosum* DSM 1402, CP036346.1 | 3.25 | *spoIIM*, *spoIIIAA, spoIIIAB, spoIIIAC, spoIIIAD, spoIIIAE, spoIIIAF, spoIIIAG, spoIIID, spoIVFA, spoVG, spoVT, spoVID, bofA, bofC, cgeD, cotQ, gerAB, gerAC, gerD, gerW/ytfJ, lipC, safA, sleL/yaaH/ydhD*, *yabQ, ymfJ, ypeB, yqfC, yrkC, yyaC, yyaD* |
| ***Limnochordia*** | | |
| *Limnochorda pilosa,* AP014924.1 | 3.82 | *spoVG, bofA, yaaT* |

**Table S5. Conservation of the previously defined sporulation core[a]**

| Sporulation stage | Phylogenetic distribution of the genes | | |
|---|---|---|---|
| | **All spore-forming bacilli and clostridia[b]** | **All bacilli and most clostridia** | **Most bacilli, some clostridia** |
| **Stage 0 (pre-septation)** | ***spo0A***, ***sigH (spo0H)***, *spo0J*, *spo0JB (yyaA)*, ***obgE*** | *spo0E*, *rapA(spo0L) family*, *ylbF*, *yjcM*, *yyaA* | *spo0M*, *spo0F*, *ytxC* |
| **Stage II (post-septation)** | ***spoIIAA***, *spoIIAB*, ***sigF (spoIIAC)***, ***spoIID***, *spoIIE (spoIIH)*, ***spoIIGA***, ***sigE (spoIIGB)***, ***spoIIM***, ***spoIIP***, ***spoIIR*** | | |
| **Stages III-VI (post-engulfment)** | ***cwlD***, ***dacB***, ***spmA***, ***spmB***, ***spoIIIAA***, ***spoIIIAB***, ***spoIIIAC***, ***spoIIIAD***, ***spoIIIAE***, ***spoIIIAF***, ***spoIIIAG***, ***spoIIIAH***, ***spoIIID***, *spoIIIE*, *spoIIIJ*, *jag*, ***sigG (spoIIIG)***, ***sigK (spoIVCB+spoIIIC)***, ***spoIVB***, *spoVAC*, *spoVAD*, *spoVAEB*, ***spoVB family***[c], ***pth (spoVC)***, ***spoVD***, ***spoVG***, ***spoVK***, ***spoVS***, *spoVT*, ***stoA (spoIVH)***, ***yabP***, ***yabQ***, ***ylbJ***, *ymxH*, ***yqfC***, ***yqfD***, *ytvI*, *yyaC* | ***bofA***, ***spoIVFB***, *spoVAEA*, *spoVAF*, ***spoVE***, ***dpaA (spoVFA)***, ***dpaB (spoVFB)***, ***ald (spoVN)***, ***spoVR***, ***sspA family***[c], *ydcC*, *yhbH*, *yqfU*, ***ytrH***, ***yunB*** | *spoVAA*, *spoVAB*, *yfhM*, *ykuD*, *ypqA*, *yqfS*, *ytrI* |
| **Spore coat** | ***spoIVA***, *alr (yncD)*, *yhaX* | ***spoVM***, *cotJC/yjqC*, *cotF* family, *lipC (ycsK)*, *yaaH*, *yabG*, *ydhD*, *yhaX*, *yhbA*, *yhbB*, *yhcN*, *yhcQ*, *yhjR* | *cotA*, *cotC*, *cotH*, *cotI*, *cotJA*, *cotJB*, *cotM*, *cotP*, *cotS*, *cotU*, *tgl*, *yisY*, *yknT* |
| **Germination** | *gpr*, *lgt (gerF)* | ***gerA family***[c], ***gerB family***[c], ***gerC family***[c], *ypeB*, *ytgP* | |

[a] – This is an updated version of Table 3 from Galperin *et al.* (2012) Environ. Microbiol. 14(11), 2870–2890). Genes that appear to be essential for sporulation of *B. subtilis* are shown in bold typeface. Gene names in parentheses indicate alternative names of the same genes.

[b] – The genes in the table are colored as follows: green, core sporulation genes (found in at least 75 out of 76 spore-formers in the analyzed genome set); yellow, widespread sporulation genes (missing in just a few spore-formers); red, genes missing in at least 10 spore formers; blue, genes conserved in bacilli (but not necessarily in clostridia); purple, genes missing in multiple bacilli and clostridia; gray, excluded from the current list because of the absence of the COG (*cotC*, *cotU*). Two genes, *rapA(spo0L)* and *yhbA*(*queG*) were excluded because of insufficient evidence for their role in sporulation; *yknT (cse15)* was excluded because of its low-complexity sequence; *ytgP* was included in the SpoVB family.

[c] – The *cotF* family includes *cotF*, *yhcQ*, *yraD*, *yraF*, and *yusN* genes; *gerA* family includes *gerAA*, *gerBA*, *gerKA*, *yfkQ*, and *yndD* genes; *gerB* family includes *gerAB*, *gerBB*, *gerKB*, *gerXB*, *yfkT*, and *yndE* genes; *gerC* family includes *gerAC*, *gerBC*, *gerKC*, *yfkR*, and *yndF* genes; *rapA* family includes *rapA*, *rapB*, *rapC*, *rapD*, *rapE*, *rapF*, *rapG*, *rapH*, *rapI*, *rapJ*, and *rapK*; *spoVB family includes spoVB*, *ykvU* and *ytgP; sspA* family includes *sspA*, *sspB*, *sspC*, and *sspD* genes

**Table S6. Core sporulation genes missing in certain genomes[a] (by gene)**

| Gene | COG no. | Missing in the following organisms[b] |
|---|---|---|
| *spoIID* | COG2385 | *Sulfobacillus acidophilus* |
| *spoIIM* | COG1300 | *Cellulosilyticum lentocellum*, *Intestinimonas butyriciproducens, Planococcaceae* |
| *spoIIP* | COG5819 | *Alicyclobacillus acidocaldarius, Thermoclostridium stercorarium* |
| *spoIIR* | COG5822 | [*Clostridium*] *innocuum* |
| *spoIIID* | COG5830 | *C. innocuum, Erysipelatoclostridium ramosum* |
| *spoIVFB* | COG1994 | *Cellulosilyticum lentocellum* |
| *spoVAC* | COG5836 | *Jeotgalibacillus malaysiensis* |
| *spoVAD* | COG5837 | *J. malaysiensis* |
| *spoVG* | COG2088 | *Carboxydothermus hydrogenoformans, I. butyriciproducens, E. ramosum, Limnochorda pilosa, Moorella thermoacetica, Pelosinus fermentans* |
| *spoVS* | COG2359 | *Anaerotignum propionicum, C. lentocellum, C. innocuum, I. butyriciproducens, Lachnoclostridium phytofermentans, Mahella australiensis, T. stercorarium* |
| *spoVT* | COG5845 | *E. ramosum* |
| *bofA* | COG5848 | *A. propionicum*, *Candidatus* Arthromitus sp., *Clostridioides difficile*, *C. innocuum, E. ramosum, I. butyriciproducens*, *L. pilosa. Paeniclostridium sordellii*, *Paenisporosarcina* sp., *S. acidophilus* |
| *cotJC/yjqC* | COG3546 | *Candidatus* Desulforudis audaxviator, *L. phytofermentans*, *J. malaysiensis*, *S. acidophilus*, *Virgibacillus halodenitrificans* |
| *dpaA, dpaB (spoVFA, spoVFB)* | COG5842, COG5843 | *Alkaliphilus metalliredigens, Clostridium acetobutylicum, Clostridium botulinum, Clostridium perfringens, Clostridium tetani, P. sordellii, Tepidanaerobacter acetatoxydans, Thermoanaerobacter italicus, Thermoanaerobacterium thermosaccharolyticum, Methylomusa anaerophila, P. fermentans, Gottschalkia acidurici, Tissierellia* sp. |
| *gerA* family | COG5901 | *C. difficile, P. sordellii* |
| *gerB* family | COG5902 | *C. difficile, C. innocuum, E. ramosum, P. sordellii* |
| *gerC* family | COG5903 | *C. difficile, C. innocuum, E. ramosum, P. sordellii* |
| *gerM* | COG5401 | *A. acidocaldarius, Ca.* Arthromitus sp., *C. acetobutylicum, P. sordellii, Planococcaceae, S. acidophilus* |
| *gpr* | COG5911 | *J. malaysiensis* |

| | | |
|---|---|---|
| *jag* | COG1847 | *Ca.* Arthromitus sp., *Kyrpidia tusciae, S. acidophilus* |
| *lipC* (*ycsK*) | COG2755 | *Ca*. Desulforudis, *C. innocuum, E. ramosum*, *Desulfofundulus kuznetsovii*, *G. acidurici*, *S. acidophilus*, *T. italicus*, *Ureibacillus thermosphaericus* |
| *ricT (yaaT)* | COG1774 | *Heliobacterium modesticaldum, L. pilosa* |
| *sleL/ydhD* | COG3858 | *Anoxybacillus flavithermus*, *C. hydrogenoformans*, *C. difficile*, *C. innocuum, C. perfringens*, *E. ramosum*, *Paenisporosarcina* sp., *P. sordellii* |
| *spmA* | COG2715 | *Planococcaceae* |
| *spmB* | COG0700 | *Planococcaceae* |
| *sspA* | COG5852 | *C. innocuum* |
| *yabQ* | COG5869 | *C. innocuum, E. ramosum* |
| *yisY* | COG0596 | *Amphibacillus xylanus*, *Ca.* Desulforudis, *Pelotomaculum thermopropionicum* |
| *yitE* | COG1284 | *Ca.* Desulforudis |
| *ylmC/ymxH* | COG1873 | *C. lentocellum* |
| *ypeB* | COG5914 | *A. propionicum*, *Ca*. Arthromitus, *C. difficile*, *C. innocuum, C. lentocellum*, *C. perfringens*, *E. ramosum*, *J. malaysiensis*, *L. phytofermentans*, *P. sordellii*, *M. anaerophila*, *P. fermentans* |
| *yqfC* | COG5871 | *E. ramosum, Planococcaceae* |
| *yyaC* | COG5878 | *A. propionicum, Ca.* Arthromitus, *C. innocuum, E. ramosum*, *I. butyriciproducens* |

[a] – Frameshifts and nonsense mutations (Table S3) are ignored. Gene names in parentheses indicate alternative names of the same genes, slash indicates paralogs (members of the same COG).

[b] – Full organism names and genome assemblies are listed in Table S1. The genus names are spelled in full at the first instance and in abbreviated form after that. "*Planococaceae*" indicates ≥5 organisms from that family.

#### Table S7. Split *sigK* genes in diverse firmicutes

| Organism name, GenBank accession number | Genomic locus tag (frame), ORF length[a] | | | Predicted *skin* element[b] | |
|---|---|---|---|---|---|
| | *spoIVCB* | *spoIIIC* | *spoIVCA* | Location | Size, ORFs |
| **Bacilli** | | | | | |
| *Bacillus subtilis* subsp. *subtilis* str. 168, AL009126.3 | BSU_25760 (+3), 129 aa | BSU_26390 (+3), 138 aa | BSU_25770 (-), 500 aa | 2,653,463 – 2,701,338 | 47.9 kb, 67 ORFs |
| *Anoxybacillus flavithermus* WK1, CP000922.1 | Aflv_0804 (-3), 129 aa | Aflv_0775 (-3), 113 aa | Aflv_0803 (+), 498 aa | 817,174 – 834,234 | 16.1 kb, 28 ORFs |
| *Brevibacillus brevis* NBRC 100599, AP008955.1 | BBR47_19200 (+3), 120 aa | BBR47_19310 (+2), 121 aa | BBR47_19300 (+), 499 aa | 2,017,400 – 2,026,645 | 9.2 kb, 10 ORFs |
| *Novibacillus thermophilus* SG-1, CP019699.1 | B0W44_05180 (+1), 119 aa | B0W44_05435 (+3), 118 aa | B0W44_05430 (+), 499 aa | 1,053,756 – 1,092,225 | 38.5 kb, 50 ORFs |
| **Clostridia** | | | | | |
| *Alkaliphilus metalliredigens* QYMF: CP000724.1 | Amet_2474 (+3), 117 aa | Amet_2480 (+2), 123 aa | Amet_2475 (-), 528 aa | 2,523,887 – 2,527,436 | 3.5 kb, 4 ORFs |
| *Caproiciproducens* sp. NJN-50, CP035283.1 | EQM14_04205 (-1), 114 aa | EQM14_04145 (-3), 114 aa | EQM14_04150 (-), 562 aa | 856,703 – 868,853 | 12.1 kb, 11 ORFs |
| *Clostridioides difficile* 630, AM180355.1 | CD630_12300 (-2), 130 aa | CD630_12300 (-3), 89 aa | CD630_12310 (-), 505 aa | 1,429,690 – 1,443,350 | 14.6 kb, 18 ORFs |
| *Clostridium tetani* E88, AE015927.1 | CTC_01066 (+3), 150 aa | N/A, 1,185,018 to 1,185,302, 95 aa | CTC_01067 (-), 502 aa | 1,138,064 – 1,185,018 | 47.0 kb, 46 ORFs |
| *Natranaerobius thermophilus* JW/NM-WN-LF, CP001034.1 | Nther_1770 (-2), 114 aa | Nther_1737 (-2), 113 aa | Nther_1738 (-), 505 aa | 1,803,258 – 1,837,025 | 33.8 kb, 30 ORFs |
| *Pelotomaculum thermopropi-onicum* SI, AP009389.1 | PTH_1126 (-3), 118 aa | PTH_1122 (-1), 118 aa | PTH_1125 (+), 502 aa | 1,160,494 – 1,164,714 | 4.2 kb, 2 ORFs |
| *Ruminiclostridium cellulolyti-cum* H10, CP001348.1 | Ccel_0434 (+2), 138 aa | Ccel_0434 (+1), 128 aa | Ccel_0440 (+), 552 aa | 498,973 – 505,588 | 6.6 kb, 6 ORFs |
| *Sulfobacillus acidophilus* TPY, CP002901.1 | TPY_3328 (-3), 139 aa | TPY_3323 (-2), 120 aa | TPY_3324 (-), 487 aa | 3,074,214 – 3,076,842 | 2.6 kb, 4 ORFs |
| *Thermacetogenium phaeum* DSM 12270, CP003732.1 | Tph_c16780 (+3), 135 aa | Tph_c16830 (+3), 129 aa | Tph_c16790 (-), 517 aa | 1,687,787 – 1,691,412 | 3.6 kb, 4 ORFs |
| **Negativicutes** | | | | | |
| *Pelosinus fermentans* JBW45, CP010978.1 | JBW_01920 (+3), 152 aa | N/A, 2233164 to 2233409, 95 aa | JBW_01921 (-), 517 aa | 2,227,052 – 2,233,164 | 6.1 kb, 5 ORFs |
| **Tissierellia** | | | | | |
| *Gottschalkia acidurici* 9a, CP003326.1 | Curi_c13480 (+3), 128 aa | Curi_c13620 (+2), 120 aa | Curi_c13490 (-), 546 aa | 1,450,043 – 1,461,566 | 11.5 kb, 13 ORFs |

[a] – The frame in the given genomic entry. ORF lengths are calculated from tBLASTn and CD-search results; GenBank entries of SpoIVCA fragments from *A. metalliredigens* (ABR48268.1), *Caproiciproducens* sp. NJN-50 (QAT49039.1), *C. tetani* (AAO35649.1), and *T. phaeum* (AFV11881.1), show longer ORFs. N/A, not annotated in the genome.

[b] – The locations and sizes of *skin* elements are calculated based on the positions of *spoIVCA* and *spoIIIC* ORFs without an adjustment for the positions of the repeat regions at the ends of the *skin* elements. The numbers of ORFs encoded in the *skin* elements are taken from the respective genomic entries.

**Table S8. COG-based annotation of uncharacterized sporulation protein**

| Gene | BSU no. | Regu lator | UniProt name | UniProt annotation | Member of COG no. | COG annotation |
|---|---|---|---|---|---|---|
| *ydgB* | BSU05570 | SigK | YDGB_BACSU | Uncharacterized protein YdgB | COG5911 | Spore germination protease Gpr |
| *ylyA* | BSU15440 | SigG | YLYA_BACSU | Uncharacterized protein YlyA | COG1734 | RNA polymerase-binding transcription factor DksA |
| *yoaR* | BSU18720 | SigG | YOAR_BACSU | Uncharacterized protein YoaR | COG2720 | Vancomycin resistance protein YoaR (function unknown), contains peptidoglycan-binding and VanW domains |
| *ypqA* | BSU22240 | SigE | YPQA_BACSU | Uncharacterized protein YpqA | COG0071 | Small heat shock protein IbpA, HSP20 family |
| *yqcK* | BSU25800 | SigF | YQCK_BACSU | Uncharacterized protein YqcK | COG0346 | Catechol 2,3-dioxygenase or related enzyme, vicinal oxygen chelate (VOC) family |
| *yrkC* | BSU26560 | SigK | YRKC_BACSU | Uncharacterized protein YrkC | COG0662 | Mannose-6-phosphate isomerase, cupin superfamily |
| *yteA* | BSU30840 | SigG | YTEA_BACSU | Uncharacterized protein YteA | COG1734 | RNA polymerase-binding transcription factor DksA |
| *ytlA* | BSU30600 | SigK | YTLA_BACSU | Putative binding protein YtlA | COG0715 | ABC-type nitrate/sulfonate/bicarbonate transport system, periplasmic component |
| *ytlC* | BSU30610 | SigK | YTLC_BACSU | Uncharacterized ABC trans-porter ATP-binding protein | COG1116 | ABC-type nitrate/sulfonate/bicarbonate transport system, ATPase component |
| *ytlD* | BSU30620 | SigK | YTLD_BACSU | Uncharacterized ABC trans-porter permease protein | COG0600 | ABC-type nitrate/sulfonate/bicarbonate transport system, permease component |
| *ywcA* | BSU38240 | SigE | YWCA_BACSU | Uncharacterized symporter YwcA | COG4147 | Na+(or H+)/acetate symporter ActP |
| *ycgL* | BSU03190 | SigE | YCGL_BACSU | Uncharacterized protein YcgL | COG3541 | Predicted nucleotidyltransferase YcgL |
| *yjaZ* | BSU11350 | SigE | YJAZ_BACSU | Uncharacterized protein YjaZ | COG5504 | Predicted Zn-dependent protease YjaZ, DUF2268 family |
| *yqhO* | BSU24510 | SigE | YQHO_BACSU | Uncharacterized protein YqhO | COG1752 | Predicted acylesterase/phospholipase RssA, containd patatin domain |
| *ylaK* | BSU14810 | SigE | YLAK_BACSU | Uncharacterized protein YlaK | COG1875 | Predicted ribonuclease YlaK, contains NYN-type RNase and PhoH-family ATPase domains |
| *yvnB* | BSU35040 | SigG | YVNB_BACSU | Uncharacterized protein YvnB | COG1409 | 3',5'-cyclic AMP phosphodiesterase CpdA |
| *yqjC* | BSU23930 | SigG | YQJC_BACSU | Uncharacterized protein YqjC | COG0346 | Catechol 2,3-dioxygenase or related enzyme, vicinal oxygen chelate (VOC) family |
